# Cerebellar growth is associated with domain-specific cerebral maturation and socio-linguistic behavioral outcomes

**DOI:** 10.1101/2025.09.11.675513

**Authors:** Aikaterina Manoli, Neville Magielse, Felix Hoffstaedter, Nilsu Sağlam, Thanos Tsigaras, Augustijn A.A. de Boer, Lorenz Ahle, Ceyda Yalçin, Milin Kim, Torgeir Moberget, Thomas Wolfers, Casey Paquola, Charlotte Grosse Wiesmann, Andre F. Marquand, Jörn Diedrichsen, Sofie L. Valk

## Abstract

The cerebellum’s involvement in cognitive functions is increasingly recognized, yet its developmental contribution to cognition remains poorly understood. The cerebellum undergoes rapid development in early life, paralleling major cognitive and behavioral changes. Although clinical studies have linked early cerebellar disruptions to profound developmental deficits, it remains largely unclear how typical cerebellar maturation supports the development of cognitive functions and how it interacts with broader brain development. Here, we apply a normative modeling framework to map cerebellar volumetric growth from infancy to young adulthood (*N* = 751; ages 1-21 years). Using lobular and functional cerebellar parcellations, we comprehensively characterize typical cerebellar development and examine how it aligns with cerebral development and behavioral outcomes. Across parcellations, posterior higher association areas consistently show steeper growth trajectories than anterior sensorimotor areas. Cerebellar and cerebral areas with similar functional roles demonstrate coordinated maturation, and volumetric growth in the posterior cerebellum relates to individual differences in socio-linguistic behaviors. These findings establish a comprehensive reference for typical cerebellar development, highlight cerebellar co-maturation with the cerebral cortex, and underscore the cerebellum’s role in supporting emerging higher cognitive functions.

## Introduction

The cerebellum’s vast computational architecture is increasingly recognized as a “universal learning system”^1–3^ that supports both sensorimotor and higher cognitive processes, including language, memory, and social cognition^4–7^. The expansive functional cerebellar repertoire is suggested to be largely mediated through extensive reciprocal connections with the cerebral cortex^8–11^. Current theoretical frameworks propose that the cerebellum encodes internal models of actions and behaviors that are transmitted to cerebral regions to facilitate automated behavioral control and optimization^1,3,12,13^. Despite substantial evidence supporting the cerebellum’s role in diverse cognitive processes, its specific role in cognitive development remains poorly characterized.

The cerebellum exhibits a protracted developmental trajectory, emerging as one of the first brain structures at approximately thirty days post-conception and being among the last to achieve maturation in the first postnatal years^14–16^. This extended developmental timeline confers particular vulnerability to environmental perturbations that may disrupt normal cerebellar maturation. Lesions in early life, when the cerebellum is still rapidly developing, result in profound and persistent cognitive impairments, which have disproportionate effects on socio-linguistic functions compared to lesions sustained in adulthood^17,18^. Moreover, cerebellar abnormalities, including volumetric reductions and atypical functional activation patterns, are consistently associated with neurodevelopmental conditions such as autism spectrum disorder (ASD), attention deficit hyperactivity disorder (ADHD), and schizophrenia^19–21^, as well as deficits across multiple cognitive and affective domains^22^. Characterizing typical cerebellar development is therefore essential for identifying critical periods during which developmental disruptions may exert substantial and enduring effects on neurodevelopmental and cognitive outcomes^23^.

Normative modeling offers a powerful framework for characterizing cerebellar developmental trajectories. By modelling developmental trajectories across many individuals, population-based normative models provide a reference that captures the expected range of typical development within a neural circuit^24,25^. Unlike traditional group-average approaches that assume homogeneity within populations, normative modeling quantifies individual deviations from typical patterns, enabling the identification of variable developmental profiles and their associations with cognitive and behavioral outcomes^26,27^. Furthermore, this framework supports robust modeling of normative development using parameters derived from large, multi-site datasets and can be flexibly updated as new data become available. Currently, a unified normative model spanning the entire cerebellar developmental trajectory from infancy through young adulthood is lacking. Recent normative modeling studies focusing on childhood and adolescence have revealed that anterior and posterior cerebellar subregions mature at different rates^28,29^. These findings are complemented by longitudinal research demonstrating rapid growth across all cerebellar lobules in the first two years of life, and especially during the first seven months after birth^30^. Despite this progress, a comprehensive reference model spanning infancy to adulthood remains critically needed, given the different rates in which the cerebellum matures in the first postnatal years compared to childhood and adolescence^14–16,18^. Addressing this gap is important, as cerebellar subregions may exhibit divergent maturation patterns across developmental stages, potentially indicating periods when different regions are most susceptible to developmental perturbations^21^. Establishing a comprehensive reference model of cerebellar growth from infancy to adulthood would thus enable more precise estimates of individual variability related to both typical and atypical cerebellar development.

Furthermore, a critical aspect of understanding cerebellar regional development involves examining its relationship with cerebral cortical maturation. The cerebellum does not act as an isolated neural circuit but interacts extensively with the cerebral cortex through numerous reciprocal pathways^31^, where it plays a fundamental role in supporting cerebral functions^1,13,32,33^. It has been proposed that cerebellar topography may largely mirror that of the cerebral cortex^34^. Consequently, functionally related regions of the cerebellum and cerebral cortex may exhibit coordinated maturation, with areas subserving similar functions displaying similar growth patterns^35^. By mapping typical cerebellar and cerebral development together, we can better understand how their coordinated growth supports integrated brain function and how variation in one system may be associated with alterations in the other.

Finally, understanding how cerebellar development contributes to cognitive and behavioral capabilities remains a key area for investigation. Functional magnetic resonance imaging (fMRI) studies^36–38^ and meta-analytic investigations^39,40^ have established detailed functional parcellations of the adult cerebellum based on its involvement in diverse motor, affective, and higher association domains. These studies reveal that anterior cerebellar regions are primarily involved in sensorimotor functions, while posterior regions, particularly Crus I and II, are engaged in higher-order cognitive functions, including language and social cognition^4,6^. However, how the cerebellum supports these functions during development remains poorly understood. Posterior cerebellar maturation is linked to early-life acquisition of several behaviors, including fine motor skills^30^ and social cognitive abilities^32^, as well as vulnerability to neurodevelopmental disorders^28,29^. This diversity of behavioral outcomes highlights the need for a comprehensive and detailed investigation of cerebellar growth trajectories and their relation to behavioral development.

To investigate cerebellar development and how it relates to that of the cerebral cortex and behavioral outcomes, the present study describes normative models of cerebellar volumetric growth from infancy to young adulthood (1-21 years). We used cross-sectional infant data (1-2 years; *N* = 111) from the Baby Connectome Project (BCP)^41^ and child and adolescent data (5-21 years; *N* = 640) from the Lifespan Human Connectome Project in Development (HCP-D)^42^. Recognizing that functional boundaries in the cerebellum do not coincide well with the anatomical, lobular boundaries^4^, we examined cerebellar development across both lobular and functional parcellations to comprehensively characterize subregional development in relation to diverse cognitive functions. For lobular boundaries, we applied a deep learning segmentation of cerebellar lobules (ACAPULCO)^43^. For functional boundaries, we leveraged multiple atlases: cerebellar resting-state networks^36^, a parcellation based on a multi-domain task battery (MDTB)^4^, and a parcellation constructed by fusing diverse task-based datasets into a single functional atlas^6^. We further associated cerebellar normative growth trajectories with developmental patterns of the cerebral cortex to examine maturational coupling between cerebellar and cerebral subregions. Finally, we investigated how subregional cerebellar growth relates to performance across a range of behavioral domains, including sensorimotor abilities, language, memory, and social cognition. Our results show that posterior cerebellar regions involved in higher association processes undergo more rapid developmental growth than anterior sensorimotor regions. We also find evidence of coordinated maturation between cerebellar and cerebral regions with similar functional roles. Notably, we observe a selective relationship between posterior cerebellar maturation and individual variation in socio-linguistic abilities. This is particularly strong for social behavior, where cerebellar development explains variance beyond what cerebral cortical development alone can account for. Together, these findings deepen our understanding of how cerebellar development aligns with cerebral maturation and supports emerging higher cognitive abilities, with potential relevance for identifying biomarkers of atypical neurodevelopment.

## Results

### Posterior cerebellar lobules exhibit steeper trajectories than anterior lobules from infancy to adulthood

We generated normative models of cerebellar anatomical lobules for ages 1-21 with Hierarchical Bayesian Regression (HBR) implemented in the PCNtoolkit^25,44,45^. Both linear and third-order B-spline models were generated. Both performed equally well based on leave-one-out cross-validation (**Supplementary Table 5**). For parsimony, we here described linear models, though B-spline models are also provided as they may offer greater flexibility for future applications^28^. Posterior distributions for all model parameters and parcels demonstrated good convergence (over 95% of R-hat values below 1.01^46^).

The lobular segmentation underwent rigorous quality control, with each image independently assessed and, if necessary, carefully manually corrected by multiple reviewers through a consensus-based procedure (see **Methods: Cerebellar parcellation**). Lobules demonstrated volumetric increases from infancy to adulthood (**Figure 1**, **Supplementary Figure 5,** and **Supplementary Table 1**), with the steepest trajectories observed in the corpus medullare (cerebellar white matter) (mean standardized age beta (*β_ag_*_e_) = 0.72; 95% confidence interval (CI) = [0.09 1.44]) and bilateral VIIB (left: *β_ag_*_e_ = 0.63; 95% CI = [0.18 1.52]; right: *β_ag_*_e_ = 0.79; 95% CI = [0.23 1.73]), followed by the right VI (*β_ag_*_e_ = 0.47; 95% CI = [-0.16 1.44]), right Crus I (*β_ag_*_e_ = 0.47; 95% CI = [-0.14 1.06]), and bilateral Crus II (left: = *β_ag_*_e_ = 0.36; 95% CI = [-0.21 1.18]; right: *β_ag_*_e_ = 0.37; 95% CI = [-0.17 1.17]). Overall, posterior lobules (VI-X: overall mean *β_ag_*_e_ = 0.29) exhibited steeper trajectories than anterior lobules (I-V: overall mean *β_ag_*_e_ = 0.25). We also observed subtle sex differences across parcellations, with age-related coefficients being slightly higher for males (*β_ag_*_e_ across parcels = 0.29) than for females (*β_ag_*_e_ across parcels = 0.28). The largest sex differences were observed in the corpus medullare (mean absolute difference in *β_ag_*_e_ = 0.52) (**Figure 1A**, bottom).

**Figure 1.**
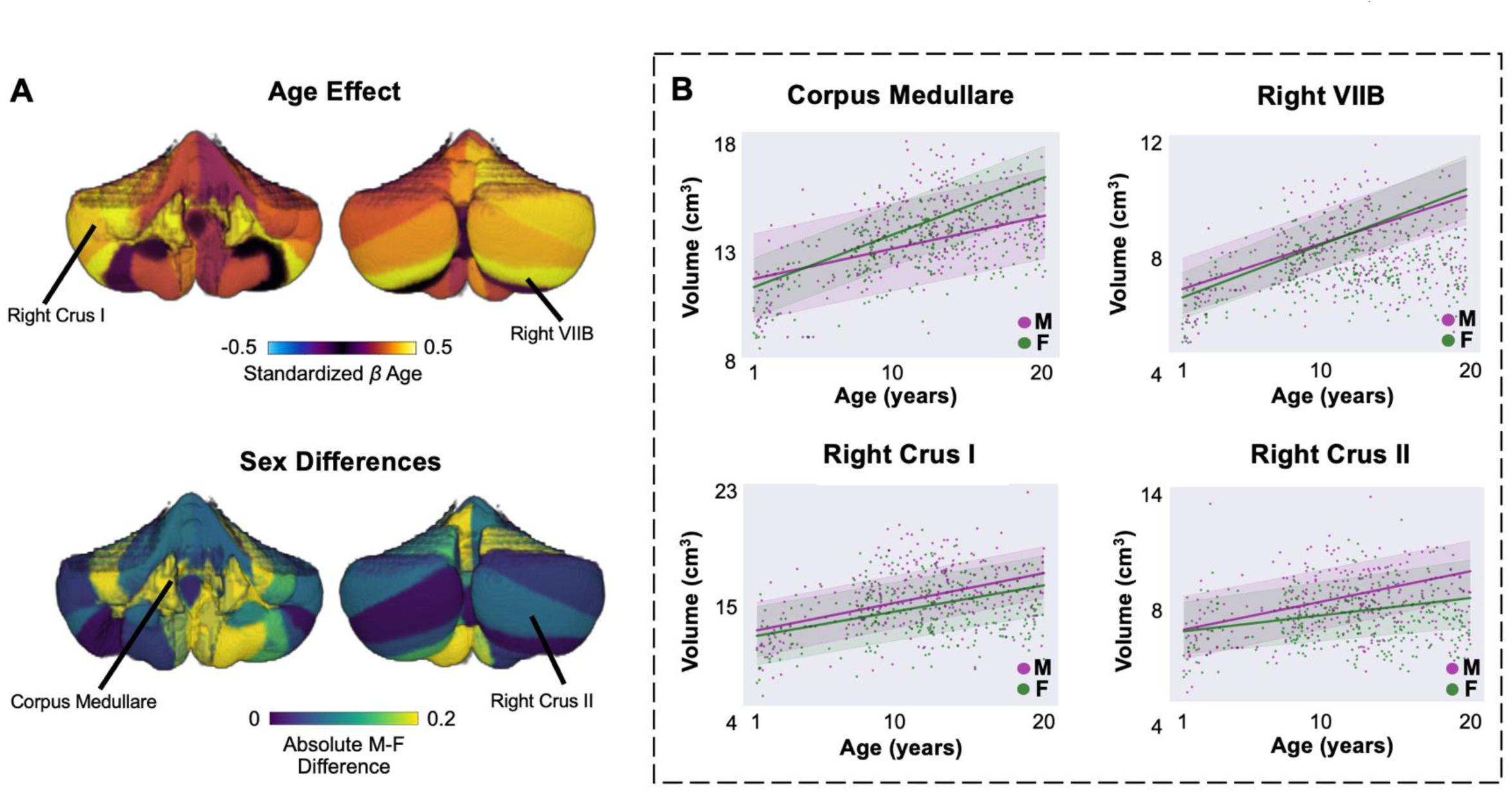
Normative growth trajectories for cerebellar anatomical lobules from infancy to adulthood (1-21 years). **A.** Mean effect of age on growth of anatomical lobules (top) and absolute male-female sex differences (bottom). Mean *β_ag_*_e_ values for anterior lobules I-IV are combined. **B.** Sex-stratified normative trajectories for four of the lobules with the steepest developmental trajectories. Bold lines represent the mean trajectories per sex. Shaded areas represent the 95.5% confidence interval. Abbreviations: M = male; F = female.

### Cerebellar higher associative regions exhibit steeper growth trajectories than sensorimotor regions from childhood to adulthood

Next, we generated normative models for three widely used functional parcellations derived from task-based and resting-state neuroimaging data^4,6,36^ to comprehensively describe normative cerebellar development across different functional taxonomies. The parcellations and associated parcel labels can be seen in detail in **Supplementary Figure 4**. Consistent with the lobular analyses, we describe results from linear models, which showed predictive performance similar to that of third-order B-spline models as assessed by LOOCV (**Supplementary Tables 6-8**).

We restricted analyses to participants aged 5 years and older, as functional spatial patterns in the first years of life may differ from those in adults but show similarity with adult functions later in childhood^47,48^. General normative developmental patterns mirrored those of the lobular trajectories and were consistent across parcellations. Specifically, we again observed age-related volumetric increases across all functional parcellations in both males and females. In all parcellations, anterior parcels, primarily associated with sensorimotor functions, exhibited smaller age-related effects and less steep growth trajectories compared to posterior parcels, associated with a range of higher-order cognitive processes (e.g., language, memory; **Figure 2A-C**). For instance, left S1 of the fusion atlas (linguistic processing) showed the highest age effect (*β_ag_*_e_ = 0.42; 95% CI = [0.32 0.52]), while left M4 (motor processing) showed the lowest (*β_ag_*_e_ = 0.08; 95% CI = [- 0.01 0.32]).

**Figure 2.**
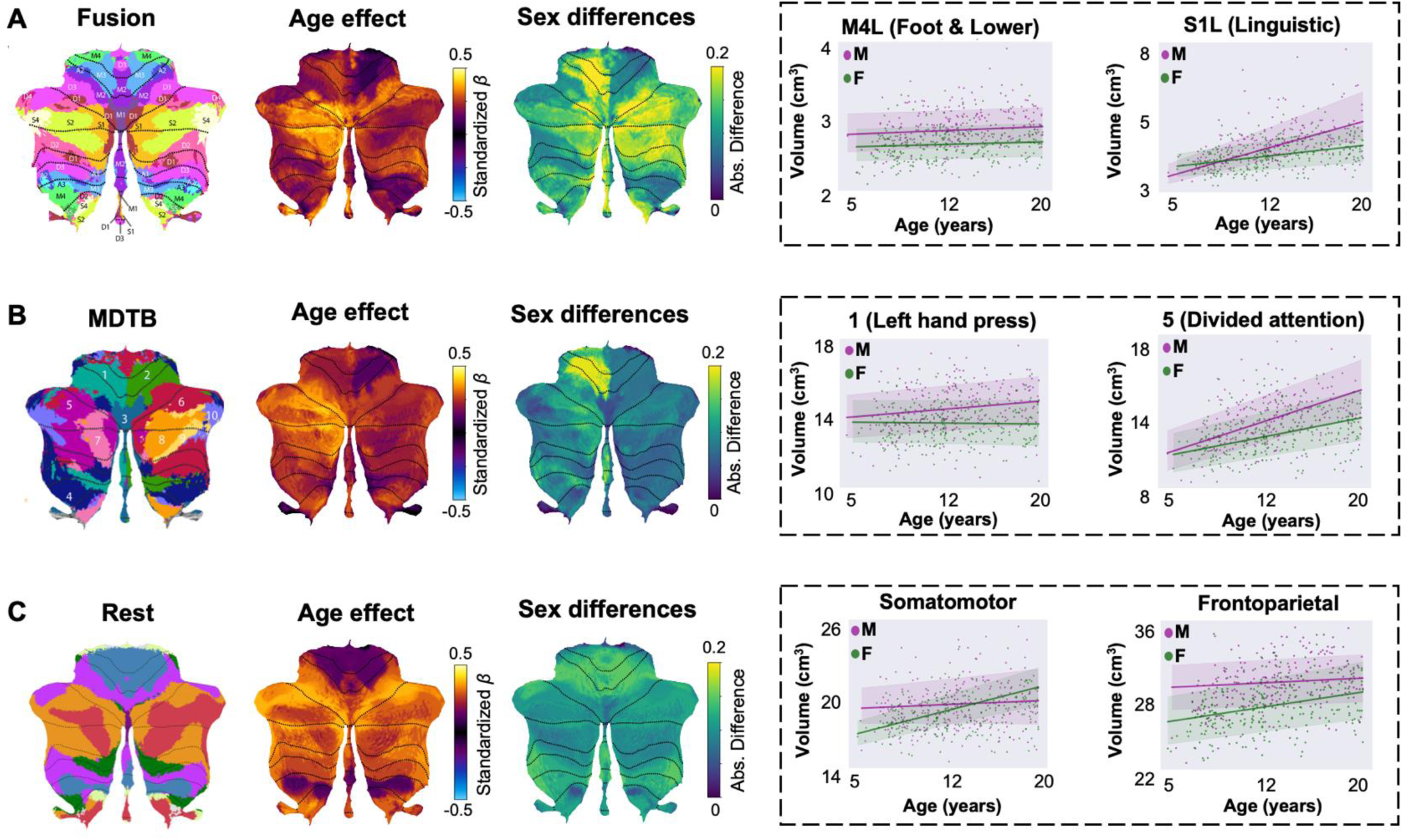
Normative growth trajectories for cerebellar functional parcels from childhood to adulthood (5-21 years). **A-C.** Mean age effects, sex differences and example trajectories for sensorimotor and higher association parcels are shown for the fusion **(A)**, MDTB **(B)**, and resting-state atlas **(C).** Left: Atlas images, adapted with permission from Nettekoven et al. (fusion)^6^, King et al. (MDTB)^4^, and Buckner et al. (rest)^36^. Middle: Mean effect of age and absolute male-female sex differences of functional lobules. Right: Sex-stratified normative trajectories for example parcels. Bold lines represent the mean trajectories per sex. Shaded areas represent the 95.5% confidence interval. Abbreviations: Abs. = absolute; M = male; F = female; L = left; R = right.

Across all parcels, we also observed sex differences, especially in the right D2 (mean absolute difference in *β_ag_*_e_ = 0.20), left M3 (mean absolute difference in *β_ag_*_e_ = 0.20), and right S2 (mean absolute difference in *β_ag_*_e_ = 0.20) in the fusion atlas and region 1 (left hand press) in the MDTB atlas (mean absolute difference in *β_ag_*_e_ = 0.20). Overall, males exhibited higher age-related coefficients than females (**Table 1**). Mean age effects and normative trajectories of all functional parcels are provided in **Supplementary** Figures 6-8 and **Supplementary Tables 2-4**.

**Table 1.** Sex-stratified means of standardized *β*_age_ across functional parcels.

| Atlas | Male (Mean [95% CI]) | Female (Mean [95% CI]) |
| --- | --- | --- |
| Fusion | 0.31 [0.19 0.45] | 0.17 [0.06 0.29] |
| MDTB | 0.31 [0.19 0.43] | 0.21 [0.12 0.32] |
| Rest | 0.28 [0.22 0.46] | 0.25 [0.14 0.34] |

### Cerebellar and cerebral subregions demonstrate domain-specific maturational coupling

Next, we examined cerebellar and cerebral co-maturation patterns based on normative parcel trajectories. Using Ridge regularized regression, we predicted *z*-scores in the Desikan-Killiany (DK) cerebral cortical atlas^49^ from *z*-scores of lobular and functional cerebellar parcels (see **Methods: Association with normative trajectories of the cerebral cortex**). Given the consistency observed among functional parcellations, we only focused on *z*-scores derived from the functional fusion atlas (5-21 years; **Figure 3A**), as a more comprehensive taxonomy of cerebellar functions resulting from multiple large-scale datasets^6^.

**Figure 3.**
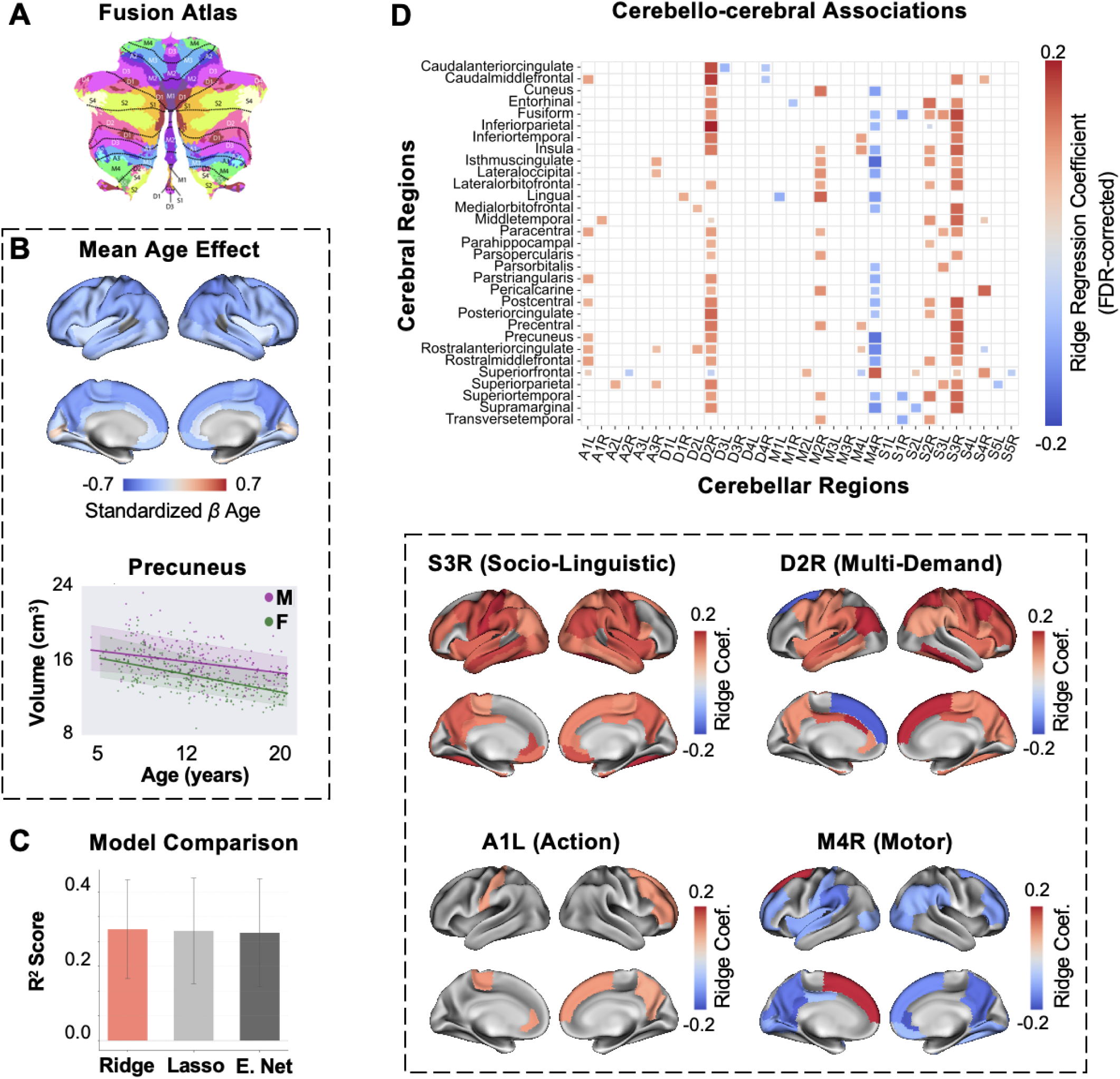
Association of cerebellar (fusion atlas) and cerebral (DK atlas) growth trajectories. **A.** Functional fusion atlas (adapted with permission from Nettekoven et al.^6^). **B.** Mean effect of age on the DK atlas (top) and example normative trajectory for precuneus (bottom). Bold lines represent the mean trajectories for each sex. Shaded areas represent the 95.5% confidence interval. **C.** Comparison of Ridge, Lasso, and ElasticNet regularization models based on average *R*^2^ scores (10-fold cross-validation). Error bars represent the standard error of the mean. **D.** Top: Significant cerebello-cerebral associations (10,000 permutations, FDR corrected at *q* = .05). Left and right DK parcels in the heatmap are averaged for brevity (see **Supplementary** Figure 10 for left and right parcel weights). Bottom: FDR-corrected weights for cerebellar parcels with the largest number of significant associations per functional domain in the fusion atlas (i.e., socio-linguistic, multi-demand, action, motor), projected on the DK atlas. Abbreviations: Coef. = coefficient; E. Net = ElasticNet; M = male; F = female; L = left; R = right.

Cerebral parcels in the DK atlas consistently exhibited volumetric decreases, with most pronounced effects observed in the precuneus (*β_ag_*_e_ = -0.23; 95% CI = [-0.60 -0.43]) (**Figure 3B** and **Supplementary Figure 9**). Cerebellar and cerebral parcels that subserved similar cognitive functions were significantly associated at the level of individual cerebellar-cerebral coefficients [10,000 permutations; one-sided and corrected for the false discovery rate (FDR)^50^ at *q* = .05], revealing domain-specific maturational coupling. The most prominent associations within each functional domain of the fusion atlas^6^ (i.e., socio-linguistic, multi-demand, motor, action) exemplify this domain-specific pattern. In the socio-linguistic domain, right S3 (corresponding to resting-state activation), was primarily associated with default-mode network regions (e.g., precuneus^51^, posterior cingulate^52^). In the multi-demand domain, right D2 (corresponding to working memory), was associated with memory and executive control regions (e.g., entorhinal^53^, caudal middle frontal gyrus^54^). Right M4 (corresponding to lower limb movements) in the motor domain was anticorrelated with several higher association cerebral regions (e.g., precuneus^51^, supramarginal gyrus^55^). Lastly, left A1 (corresponding to spatial simulation, such as mental imagery and navigation) in the action domain was linked to sensory and perceptual regions (e.g., paracentral ^55^, postcentral gyrus^57^) (**Figure 3D** and **Supplementary Figure 10**). In addition to associations with the functional fusion atlas, we also examined the relationship of lobular normative trajectories with cerebral development. This revealed similar maturational coupling patterns, particularly in posterior lobules, whose growth trajectories were associated with cerebral higher association regions. However, these findings demonstrated less domain specificity, which could be explained by the observation that multiple functional processes map to each cerebellar lobule^4^ (**Supplementary** Figure 11). Overall, cerebello-cerebral maturational coupling followed functional network organization, with functionally related regions exhibiting coordinated developmental trajectories across brain structures.

### Growth of cerebellar higher associative regions corresponds to socio-linguistic behavioral outcomes

Lastly, we used Partial Least Squares (PLS) analysis to link individual deviations from normative cerebellar development with individual differences in a broad range of behavioral markers. PLS identifies latent variables that capture shared patterns of variance between parcel-level *z*-scores and behavioral outcomes^58^. We applied this approach to examine associations between normative *z*-scores across fusion atlas parcels (ages 5-21; HCP-D dataset) and performance on sensorimotor and higher association tasks (see **Methods: Behavioral tasks**). Behavioral outcomes were standardized and residualized for age and sex to be consistent with the parcel *z*-scores. PLS analysis extracted a significant latent variable relating parcel growth and behavioral scores (*p* = .010, one-sided). This single latent variable accounted for 78% of the covariance across the two variables (**Figure 5A; top**). Next, the PLS model was evaluated using 10-fold cross-validation (**Figure 4A; bottom)**. The mean training set correlation across folds was *r*(113) = .31, and the mean test set correlation was *r*(11) = .27. The empirical correlation between behavioral scores and parcel *z*-scores was *r*(128) = .30, *p* = .001 (one-sided and tested against a null model assuming no brain-behavior relationships over 10,000 permutations; **Figure 4B**).

**Figure 4.**
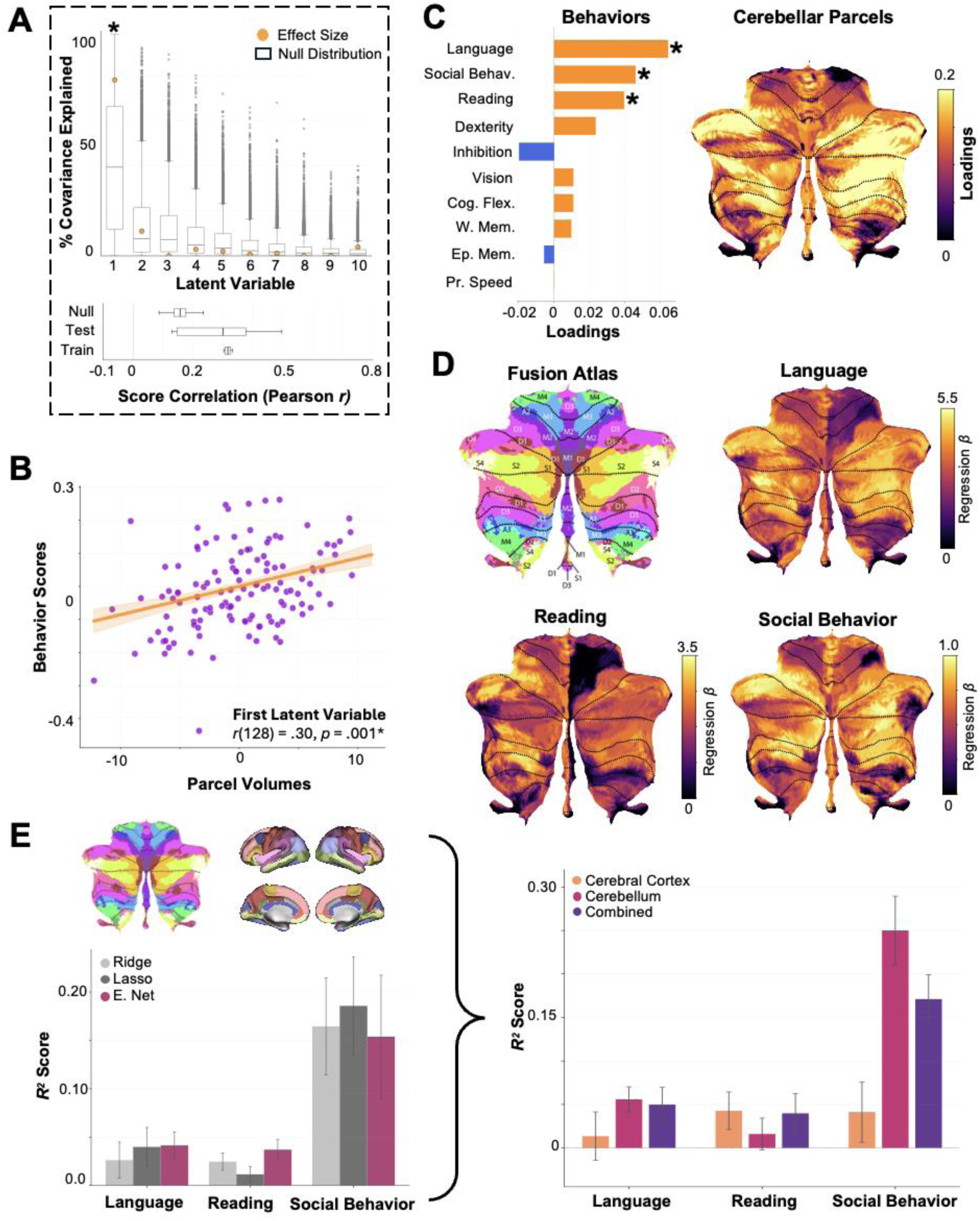
Association of cerebellar parcel-wise cerebellar growth (fusion atlas) and behavioral outcomes. **A.** Top: PLS analysis revealed a significant latent variable capturing 78% of the covariation between parcel-wise normative *z*-scores and behavioral scores (*p* = .010*, 10,000 permutations, one-sided). Bottom: We employed 10-fold cross-validation on the PLS model and assessed the statistical significance of the held-out sample correlation by comparing it to a permuted null distribution assuming no cerebellum-behavior relationships (*p* = .001*, 10,000 permutations, one-sided). **B.** Empirical correlation for the first PLS component demonstrated a significant positive relationship between cerebellar parcel scores and behavior scores (*p* = .001*, 10,000 permutations, one-sided). Data points signify individual subject scores. The line signifies the Pearson correlation coefficient, and the ribbon the 95% CI. **C**. Cerebellar parcel-wise and behavioral loadings onto the latent variable. Asterisks signify stability of behavioral outcomes (CIs not crossing zero) during 10,000 iterations of bootstrap resampling. All parcel loadings were stable during bootstrapping. Parcel loadings are projected on the cerebellar flatmap. **D.** Parcel-wise *z*-scores and standardized age and sex were used to predict behavioral outcomes in mass-univariate linear regression. This analysis was performed to map developmental behavioral associations directly onto cerebellar parcels. Only significant associations (regression betas) are plotted (*p* < .05, two-sided). **E.** Cerebellar (fusion), cerebral (Desikan-Killiany), and combined cerebello-cerebral parcel *z*-scores were compared for how well they predicted socio-linguistic behaviors using ElasticNet regularization. Left: Mean Lasso, Ridge, and ElasticNet model performance (*R^2^*) across cerebellar, cerebral, and combined models for each behavior, averaged over ten cross-validation folds. Error bars represent the standard error of the mean. Right: ElasticNet model performance (*R^2^*) for cerebellum-only, cerebral-cortex-only, and combined model across language, reading abilities and social behavior, averaged over ten cross-validation folds. Error bars represent the standard error of the mean. The functional fusion atlas was adapted with permission from Nettekoven et al., 2024. Abbreviations: E. Net = ElasticNet; Social Behav. = Social Behavior; Cog. Flex. = Cognitive Flexibility; W. Mem. = Working Memory; Ep. Mem. = Episodic Memory; Pr. Speed = Processing Speed; L = left; R = right.

The PLS analysis revealed that language comprehension, social behavior, and reading abilities had the highest loadings among behaviors, while multi-demand (e.g., D2R, D2L) and socio-linguistic (e.g., S2R, S4R) cerebellar parcels showed the highest loadings among parcels. This pattern indicates that the latent variable primarily captured covariation between socio-linguistic behaviors and higher association cerebellar regions. Notably, cerebellar regions associated with action and sensorimotor functions (e.g., A3, M4) exhibited low loadings, suggesting limited covariation with socio-linguistic behaviors. Reading, language comprehension, and social behavior demonstrated significant loadings with 95% CIs not crossing zero after 10,000 bootstrap iterations (**Figure 5C; Supplementary Figure 13**). All cerebellar parcels showed significant loadings during bootstrapping (**Supplementary Figure 13**). However, multi-demand and socio-linguistic parcels exhibited the highest bootstrap ratios (BSR = variable mean loading / standard error across iterations), indicating both strong covariation with socio-linguistic behaviors and high reliability^59^ (**Supplementary Table 9**). PLS results for socio-linguistic behaviors were further supported by mass-univariate linear regression that used parcel *z*-scores and standardized age and sex to predict behavioral scores. This analysis implicated primarily higher association cerebellum regions (social and multi-demand regions in the fusion atlas) in individual differences in socio-linguistic outcomes over development (**Figure 4D**). However, it is important to note that this analysis was performed post-hoc to decompose the PLS findings at the individual parcel level, since PLS does not identify specific univariate parcel-behavior associations^60^.

Finally, we evaluated the relative contributions of cerebellar versus cerebral normative development to socio-linguistic behavioral outcomes using ElasticNet regularized regression (see **Methods: Association with behavioral outcomes**). We compared how well normative *z*-scores from cerebellum-only, cerebral cortex-only, and combined cerebello-cerebral models predicted individual variations in language comprehension, social behavior, and reading abilities, using performance metrics (*R^2^* score) of ElasticNet models across 10 cross-validation folds (90% training, 10% testing). Cerebellum-only models performed comparably to cerebral cortex-only and combined cerebello-cerebral models for language comprehension and reading abilities (pairwise Wilcoxon signed-rank tests, all *p* > .05, two-sided; **Table 2**). However, for social behavior measured via the Social Responsiveness Scale^31^ (SRS; higher scores indicating higher autistic traits), the cerebellum-only model significantly outperformed both the cerebral cortex-only model (*W* = 0.0, *N* = 10, *p* = .002, two-sided) and the combined model (*W* = 8.0, *N* = 10, *p* = .049, two-sided) (**Figure 4E** and **Table 2**). This suggests a specific association between individual differences in normative cerebellar development and autistic traits across the developmental span from childhood to adulthood, over and above the cerebral cortex.

**Table 2.** Comparison of ElasticNet performance for cerebellum vs. other models.

| **Behavior** | **Comparison** | $M_{Cerebellum}$ | $SD_{Cerebellum}$ | $M_{Other}$ | $SD_{Other}$ | $W$ | $N$ | $p$ |
| --- | --- | --- | --- | --- | --- | --- | --- | --- |
| **Language Comprehension** | Cerebellum vs. Cerebral Cortex | 0.06 | 0.05 | 0.01 | 0.09 | 21.00 | 10 | .557 |
|  | Cerebellum vs. Combined |  |  | 0.06 | 0.07 | 21.00 | 10 | .770 |
| **Reading** | Cerebellum vs. Cerebral Cortex | 0.02 | 0.06 | 0.05 | 0.08 | 11.00 | 10 | .084 |
|  | Cerebellum vs. Combined |  |  | 0.04 | 0.07 | 10.00 | 10 | .083 |
| **Social Behavior (SRS)** | Cerebellum vs. Cerebral Cortex | 0.24 | 0.13 | 0.03 | 0.13 | 0.00 | 10 | .002* |
|  | Cerebellum vs. Combined |  |  | 0.18 | 0.11 | 8.00 | 10 | .049* |

\* = $p < .05$ , two-sided

Together, these findings indicate that normative variation in cerebellar development, particularly within posterior higher associative regions, is related to individual differences in socio-linguistic behaviors. While model performance for language comprehension and reading was comparable across cerebellar and cerebral cortical (or combined) models, cerebellar parcels outperformed the other models in prediction of (atypical) social behavior. This converging evidence across multivariate and univariate approaches supports a selective relationship between posterior cerebellar maturation and individual variation in socio-linguistic abilities, and particularly the prediction of social behavior.

## Discussion

The present study describes a comprehensive normative framework for cerebellar development from the first years of life through adulthood. Across lobular, task-based, and resting-state parcellations, posterior higher association cerebellar regions demonstrated steeper growth trajectories compared to anterior sensorimotor regions. Cerebellar maturation was associated with functionally similar cerebral regions, with cerebellar higher associative areas linked to cerebral higher associative areas, and sensorimotor areas linked to cerebral sensorimotor regions, indicating functional specificity. Furthermore, developmental changes in posterior cerebellar regions associated with socio-linguistic and multiple-demand functions predicted individual differences in socio-linguistic abilities. This relationship was especially strong for social behavior, where cerebellar maturation provided predictive value beyond that of the cerebral cortex, demonstrating the cerebellum’s unique contribution to individual differences in social abilities.

All parcels, lobular and functional, exhibited age-related volumetric increases, as expected for cerebellar growth during childhood. Male participants demonstrated greater age-related effects across parcellations. This is in line with reports of sex differences in cerebellar development^28,30,61,62^ and parallels patterns observed in cerebral cortical development^24,25^. The consistent anterior-posterior differences in growth patterns across both lobular and functional parcellations mirror known patterns of cerebellar evolutionary expansion^63,64^ and significantly extend previous cerebellar developmental research^28–30,65^ in two key ways: first, by establishing a comprehensive reference of cerebellar growth spanning the entire developmental period from infancy to adulthood, and second, by systematically validating growth patterns across multiple cerebellar atlases. Steep developmental growth trajectories in posterior higher association regions across parcellations could reflect age-related improvements in underlying cognitive functions commonly associated with these regions, suggesting a contribution of the cerebellum to the development of these processes^4,6,66^. Interestingly, the lobular parcellation revealed systematic growth across both posterior and anterior cerebellar regions, particularly in anterior lobule IV. This widespread pattern reflects the extensive structural changes characterizing early postnatal cerebellar maturation^67^, extending beyond posterior associative regions alone. The developmental specificity of these changes is evident in lobular normative models in the HCP-D dataset (excluding 0-5-year-olds from BCP), where maturation was restricted to posterior regions (**Supplementary** Figure 11A). This contrast suggests that rapid anterior-posterior developmental changes occur predominantly during the first postnatal years, consistent with longitudinal studies showing accelerated growth in both anterior and posterior lobules during this period^30^. These developmental trajectories establish critical baselines for typical cerebellar development and might contribute to the detection of neurodevelopmental disorders at various developmental stages.

The pronounced influence of age on higher association posterior cerebellar regions may suggest that the cognitive functions these areas support undergo substantial refinement during childhood. Because these regions show greater age-related changes relative to anterior sensorimotor regions, they may also be more susceptible to developmental perturbations linked to neurodevelopmental or psychiatric conditions^4,15,21,67^. This is in line with findings showing that cerebellar lesions sustained in infancy are associated with more severe and lasting neurocognitive impairments than those occurring later in life^18^, emphasizing the critical role of intact cerebellar development in supporting optimal behavioral and cognitive outcomes. Our normative models, especially when integrated with clinical data, may help identify sensitive periods for the development of different cognitive functions, thus informing early intervention strategies in at-risk pediatric populations^25^.

Cerebellar parcels exhibited maturational covariance with cerebral regions implicated in the same functions. For instance, cerebellar regions associated with default-mode and social processing displayed strong associations with corresponding cerebral cortical areas (e.g., precuneus, posterior cingulate). Conversely, cerebellar regions subserving sensory functions (e.g., spatial processing) showed coordinated development with sensory cerebral areas (e.g., paracentral and postcentral gyri). These findings extend previous cerebello-cerebral covariance patterns observed in adults^35^ by demonstrating domain-specific co-maturation during childhood. This aligns with established theories of *developmental diaschisis*, wherein the cerebellum critically influences cerebral maturation^21^ through extensive reciprocal connectivity with the cerebral cortex^8–11^. Disruption of cerebellar function during critical developmental windows may consequently impair the maturation and function of connected cerebral regions^67–69^. Within this framework, our findings lay the foundations for future research to identify specific cerebral regions vulnerable to dysfunction when particular cerebellar regions are disrupted during development, based on their coordinated maturation patterns.

Furthermore, developmental variation in cerebellar circuits appears linked to individual differences in cognitive and social outcomes. We found that normative growth trajectories in posterior cerebellar regions, particularly functional subdivisions of the posterior-lateral Crus I-II, were associated with behavioral performance in language comprehension, reading, and social behavior. These findings align with growing evidence that the cerebellum, and Crus I-II in particular, plays a central role in non-motor functions such as language^70,71^ and social cognition^7,72,73^, including during development^18,32^. Additionally, both cerebellar and cerebral parcels contributed to socio-linguistic behavioral variance. Growth patterns in posterior cerebellar regions showed a particularly strong association with individual differences in social functioning as measured by the Social Responsiveness Scale^31^, which exceeded associations with the cerebral cortex. This pattern resonates with the well-documented involvement of the cerebellum in ASD neurobiology^17,21,68,74,75^ and converges with recent normative modeling work linking structural anomalies in Crus I-II to ASD traits^28,29^. Together, these results suggest that structural maturation of the posterior cerebellum is not only integral to the typical development of complex socio-cognitive functions but may also serve as a sensitive biomarker of ASD-related phenotypes, potentially even more so than the cerebral cortex. These findings reinforce the cerebellum’s emerging role as a critical contributor to higher cognitive development and highlight its relevance for early identification and monitoring of atypical social trajectories.

Our study also has important limitations. First, our sample size is on the smaller end of population-wide normative modeling endeavors. This is partly attributable to the fact that, to date, there is no reliable cerebellar segmentation method that covers the entire developmental spectrum from the perinatal period to adulthood. Different algorithms (e.g., SUIT^76,77^, ACAPULCO^43^) perform variably across age ranges, requiring laborious manual correction. Second, minimal age overlap between HCP-D and BCP datasets means age effects may be confounded with site effects, potentially limiting generalizability. Despite these constraints, our findings use careful quality control and manual correction of cerebellar segmentations to provide a comprehensive framework for cerebellar development from the first year of life through adulthood, bridging previous studies that examined infant^30^ and child/adolescent populations^28,29^ separately. Our publicly available normative models can be further validated and expanded as new data becomes available^25,28^. Future efforts should develop reliable cerebellar functional parcellation methods spanning the entire developmental trajectory, which will enable more robust normative modeling and accurate assessment of individual deviations across all developmental stages.

A further limitation is our application of adult-based functional cerebellar parcellations to developing populations. Even though children and adults may recruit similar cerebellar regions for cognitive functions (e.g., Theory of Mind^32^), functional boundaries in the cerebellum likely vary as a function of age^32^. Here, we restricted our functional atlas modeling to children and adolescents between 5-21 years. From this age onward, functional similarity to adult cerebellar organization is considered sufficient to support the use of adult-based parcellations^47,48^. However, this approach assumes that functional organization in the cerebellum remains stable across development, which likely does not fully capture age-related changes in cerebellar functional topography^78^. Future research should create age-specific functional atlases of the cerebellum across different functional tasks and developmental stages, allowing for more developmentally appropriate parcellation schemes that could reveal age-dependent spatial changes in cerebellar functional organization.

Lastly, in this study we focused on providing normative trajectories in a typically developing cross-sectional cohort. Future research should validate our models in longitudinal or clinical populations to assess individual variability over time^79^ or in or in neurodevelopmental and psychiatric conditions. For example, comparing cerebellar subregional volumes for ASD patients to our reference model could reveal its utility in ASD diagnosis. In the same vein, it can be assessed whether cerebellar or cerebello-cerebral deviations from the reference model predict patients’ behavioral outcomes. At later stages, one may use prognostic brain scans in young children to predict increased risk for several disorders or behavioral deficits. Such applications could enable early detection of cerebellar abnormalities and inform targeted interventions. The ability to quantify individual phenotypes relative to age-matched normative reference ranges represents a crucial step toward precision medicine approaches in pediatric neurology and psychiatry, particularly for conditions where cerebellar involvement is suspected or established^25^.

Overall, this study establishes normative references for human cerebellar development, providing critical reference benchmarks for detecting atypical cerebellar maturation from infancy through adulthood. Our results reveal that cerebellar posterior higher association regions systematically demonstrate prolonged growth trajectories that reflect their essential contributions to higher-order cognitive and especially social functions. The domain-specific coordination between cerebellar and cerebral maturation, combined with the predictive capacity of cerebellar volumes for social cognitive and linguistic outcomes, highlights the cerebellum’s fundamental role in neurodevelopment and its heightened susceptibility in ASD. By characterizing the relationship between cerebello-cerebral structural co-maturation and behavioral outcomes, our work deepens understanding of cerebellar development relative to cerebral and behavioral development. These findings provide a foundation for precision medicine approaches in pediatric neuropsychiatric conditions involving cerebellar dysfunction.

## Methods

### Participants

We leveraged openly available cross-sectional MRI and behavioral data of typically developing infants, children, and adolescents from the BCP^41^ and the HCP-D^42^ datasets. BCP contains structural and functional MRI data of 398 healthy term-born infants from birth to five years of age. HCP-D contains structural and functional MRI data of 652 healthy children and adolescents from five to twenty-one years of age. Informed consent was obtained from adult participants or the parents of underage participants before participation. Exclusion criteria include preterm birth, complications at birth, and medical, genetic, neurodevelopmental, or endocrine conditions. A detailed list of inclusion and exclusion criteria is provided in refs.^41,42^. In the present study, we conducted our analyses starting from the first year of life as the cutoff point. In these scans, cerebellar lobular folds were visible and image processing and segmentation algorithms performed reasonably well (**Supplementary** Figure 3), verified by visual inspection among three independent reviewers per scan (AM, NM, LA), and supervised by the project’s Principal Investigator (SLV). Overall, we excluded 94 BCP and 12 HCP-D participants due to poor image quality, insufficient cerebellar coverage, or poor performance of the segmentation algorithms after visual inspection among the same three independent reviewers. We further excluded 11 participants due to artifacts in cerebral cortical surface reconstruction (see **Cerebral cortex parcellation)** This resulted in a final sample of 751 participants (age: *M* = 12.86, *SD* = 5.49; 397 female): 111 participants from BCP (age: *M* = 2.93, *SD* = 1.36; 53 female) and 640 participants from HCP-D (age: *M* = 14.44, *SD* = 4.06; 344 female).

### Data acquisition

All data were acquired with 3T Siemens Prisma MRI scanners (Siemens, Erlangen, Germany) with 32 channel head coils. Younger participants in the BCP were scanned while naturally asleep without the use of sedatives. T1-weighted (T1w) images were acquired with a 3D MPRAGE sequence with the following parameters: sagittal field of view of 256 × 240 × 166 mm with a matrix size of 320 × 300 × 208 slices, resolution of 0.8 mm isotropic voxels, and flip angle of 8 degrees. The TR/TE parameters for BCP and HCP-D T1w images are 2400/2.24 ms and 2500/2.22 ms, respectively. Full acquisition parameters and scanning procedures are described in refs.^41,80^.

### Behavioral tasks

We examined associations between normative volumetric growth in each parcel of the functional fusion atlas and task performance in the HCP-D dataset, which provides participant scores across multiple behavioral domains. We focused on tasks that were representative of diverse sensorimotor and higher association domains and optimized for the entire HCP-D age range (5-21 years): i) the Pegboard Test for fine motor dexterity; ii) the Visual Acuity Test for visual processing; iii) the Picture Sequence Memory Task for episodic memory; iv) the List Sorting Working Memory Test for working memory; v) the Dimensional Change Card Sort Task for cognitive flexibility; vi) the Flanker Inhibitory Control and Attention Test for executive function and inhibition; vii) the Pattern Comparison Processing Speed Test for processing speed; viii) the Oral Reading Recognition Test for reading ability; ix) the Picture Vocabulary Test for language comprehension; and x) the Social Responsiveness Scale for social behavior^31^. Most of these assessments (excluding the Social Responsiveness Scale) are components of the National Institutes of Health (NIH) Toolbox (nihtoolbox.org). Detailed descriptions of all tasks are available at: humanconnectome.org/storage/app/media/documentation/LS2.0/LS_2.0_Release_Appendix_2.pdf. Distributions of participant scores for each behavioral task can be found in **Supplementary Figure 12**.

### Image preprocessing

All processing started with T1w images in participants’ native space. HCP-D images were already preprocessed according to the HCP minimal preprocessing pipeline^81^. Specifically, we utilized native space T1w images that had been aligned to T2w images and undergone bias field correction. For BCP images, we adopted a modified approach, given that these data are not currently provided in preprocessed format. BCP preprocessing steps were chosen to be consistent with the HCP-D preprocessing pipeline. For each BCP participant, one of three independent reviewers (AM, NM,

LA) conducted visual quality assessment of all scans, evaluating imaging artifacts, motion artifacts, and cerebellar coverage. Each individual-reviewer assessment was subsequently examined by the other two reviewers for consensus validation. 10% of images was then visually inspected by the project’s Principal Investigator (SLV) to ensure further validation. Quality-checked BCP images were then minimally preprocessed using iBEAT V2.0^82^, a state-of-the-art pipeline specifically optimized for infant and toddler MRI data. T1w and T2w images were reoriented to a consistent left-posterior-inferior (LPI) orientation, underwent bias field correction, and were subsequently aligned to each other. Preprocessed T1w images were then submitted to separate pipelines for cerebellar and cerebral parcellation.

### Cerebellar parcellation

Given that lobular and functional boundaries within the cerebellum do not well coincide^4^, we parcellated the cerebellum based on both lobular and functional cerebellar atlases. For the lobular segmentation, we used ACAPULCO, a validated method that uses convolutional neural networks (CNNs) to parcellate the cerebellum into its anatomical lobules^43^. The ACAPULCO algorithm was used to segment cerebella in the entire age range (1-21 years; see **Supplementary Figure 3** for example ACAPULCO segmentations in very young children in the BCP data). After registering the T1w images to the 1 mm isotropic MNI ICBM 2009c template^83^, ACAPULCO used a locating CNN to detect and fit a bounding box around the cerebellum. The cropped cerebellum was then used as input to a second parcellating CNN, which divided the cerebellum into 28 bilateral lobules. Both CNNs were trained based on expert lobular labelling in an adult (*N* = 15) and pediatric (*N* = 20) cohort (see Han et al., 2020 for details). Each MNI-based parcellation was then transformed back to original participant space via nearest neighbor interpolation. Each parcellation output underwent visual inspection and potentially manual correction for cerebellar over- and under-inclusions by three independent reviewers (AM, NM, CY) using ITK-SNAP (itksnap.org/pmwiki/pmwiki.php)^84^ (see **Supplementary Figure 2** for examples of over- and under-inclusions). To minimize potential bias, each reviewer’s corrections were subsequently examined by the other reviewers for consensus validation. 10% of manually corrected images was also visually inspected by the project’s Principal Investigator (SLV) to ensure further validation.

For the functional parcellations, we leveraged resting-state^36^ and task-based^4^ atlases of the cerebellum, as well as a recently developed atlas that fuses large-scale task-based and resting-state data from multiple datasets^6^. The resting-state atlas parcellated the cerebellum into known resting-state networks (limbic, somatomotor, ventral attention, dorsal attention, frontoparietal, and default-mode), based on resting-state functional connectivity with the same networks in the cerebral cortex^36^. The task-based (MDTB^4^) atlas used fMRI task contrast from 26 tasks (47 unique conditions) to parcellate the cerebellum. Here we used the MDTB atlas with ten functional regions: (1: Left-hand (motor) presses, 2: Right-hand (motor) presses, 3: Saccades, 4: Action observation, 5: Divided attention (left hemisphere), 6: Divided attention (right hemisphere), 7: Narrative, 8: Word comprehension, 9: Verbal fluency, 10: Autobiographical recall). Lastly, the fusion atlas^6^ parcellated the cerebellum into subregions corresponding to four large-scale functional domains (motor, action, multi-demand, socio-linguistic), derived by combining multiple datasets under a Hierarchical Bayesian framework. Here, we used the symmetric, mid-granularity version of the atlas with each of 32 bilateral regions belonging to one of the four functional domains.

Given the absence of an established functional atlas for the developing cerebellum, we applied functional atlases exclusively to the HCP-D dataset (ages 5-21 years). We did not extrapolate the adult-derived functional atlases to 1-5-year-olds, as functional spatial patterns in the first years of life may differ from those in adults^47,48^. However, such functional patterns appear to become more comparable to those of adults later in childhood, both in terms of task-specific and resting-state activity^47,48^. We calculated cerebellar volumes by resampling MNI-aligned atlases to each participant’s native space. This was achieved using nonlinear warp fields generated during spatial normalization (via FNIRT) within FSL’s (fsl.fmrib.ox.ac.uk/fsl/docs/#/) *applywarp* tool, with nearest-neighbor interpolation to preserve the discrete nature of atlas labels. We performed additional quality control by excluding participants for whom 50% or more parcels per segmentation were considered outliers (parcel volumes > |2| standard deviations of the mean volume of each parcel). Thus, we ended up excluding 13 participants for the resting-state atlas, 21 for the MDTB atlas, and 3 for the fusion atlas.

### Cerebral cortex parcellation

We obtained participant-specific cerebral cortex parcellations using FreeSurfer’s (<u>surfer.nmr.mgh.harvard.edu</u>) *recon-all* pipeline to measure the grey matter volume of parcels in the DK atlas^49^. The *recon-all* pipeline includes intensity normalization, skull stripping, tessellation of the gray and white matter cerebral boundary, automated topology correction^85^, and surface reconstruction. Surfaces were visually inspected and, when necessary, manually corrected as recommended in the FreeSurfer pipeline (<u>surfer.nmr.mgh.harvard.edu/fswiki/FsTutorial/TroubleshootingData</u>). Corrected surfaces were then registered to a spherical surface atlas and underwent gyral labelling based on the DK atlas. Gray matter volumes were calculated from the *aparc+aseg* segmentation, where each voxel is classified as belonging to a specific parcel based on probabilistic atlas information and local intensity values. Parcel volumes are computed as the sum of voxels assigned to each DK atlas region multiplied by the voxel volume. We used the Euler index (EI) to perform additional quality control of surface reconstruction. The EI was defined as the total number of surface “holes” or topological defects across both hemispheres in the cerebral surface reconstruction prior to FreeSurfer’s topological correction^24^. Given that the EI does not have a single threshold that is expected to be generalizable across different datasets^86^, in this study we excluded scans with an EI > |2| median deviations from the dataset-specific median EI^24^. This resulted in the exclusion of 11 participants (see **Participants**), who were discarded from subsequent normative modeling analyses.

### Normative modeling

We generated normative models for lobular and functional cerebellar and cerebral cortical subregions using HBR as implemented in the PCNtoolkit version 0.29^25,44,45^ in Python 3.11.8. In this way, we were able to obtain normative ranges of cerebellar volumetric growth and model individual heterogeneity (defined as deviation from the normative volumetric trajectory) with possibly substantial behavioral and clinical implications^25,45,87^. Importantly, the HBR framework allows controlling for possible confounds due to pooling data from different acquisition sites by modelling them as (random) batch effects. HBR leverages shared priors, enabling the estimation of site-specific parameters and hyperparameters. These hyperparameters can be transferred to new sites without needing the original data^26,28^, which also allows models to grow as new data becomes available. We estimated normative volumetric trajectories for each cerebellar and cerebral parcel, modelling age as the main predictor of interest and sex and acquisition site as batch effects (see **Supplementary Figure 1** for the distribution of participant ages across sites). We split our data into a training set (80%) and a test set (20%) while stratifying our sampling to ensure that sex and site were proportional among the training and test sets.

We generated linear and third-order B-spline models with five evenly spaced knot points for every parcel. We evaluated out-of-sample predictive performance of both model types using LOOCV with Pareto-smoothed importance sampling. Performance was summarized as the expected log pointwise predictive density (ELPD-LOO). Higher (less negative) values indicate better predictive accuracy. We accommodated skewed distributions through the sinh-arcsinh likelihood (SHASHb)^44^ and modeled random effects in intercept, slope, and variance (sigma) on the batch-effects (sex and site). Lastly, we used four Markov chain Monte Carlo chains with 2,000 samples, with the first 500 considered tuning samples and removed from further analyses. We included the same participants in both cerebellar and cerebral normative models to enable associations of cerebellar and cerebral developmental trajectories at the individual level.

### Association with normative trajectories of the cerebral cortex

We examined associations between individual volumetric growth trajectories (normative model-derived participant *z*-scores) of cerebellar (lobular and functional atlases) and cerebral parcels (DK atlas) to investigate maturational coupling. For the functional parcellation analysis, we focused exclusively on the functional fusion atlas^6^ given the consistent normative trajectories observed across all functional parcellations. For the lobular parcellation analysis, we restricted our investigation to the HCP-D dataset to derive accurate cerebral parcellations using FreeSurfer (FreeSurfer is not recommended for younger age ranges included in the BCP dataset^88^). This also maintained consistency with the age range employed in the functional fusion cerebellar-cerebral comparison. As explained previously, we did not extrapolate functional atlases derived in adults to individuals younger than five, to prevent unjustly projecting adult brain functions onto early childhood brains^47,48^.

We employed regularized regression to examine associations between the normative model *z*-scores of cerebellar and cerebral parcels, predicting cerebral *z*-scores from cerebellar *z*-scores. Regularization introduces penalty terms to the standard regression loss function, placing constraints on model coefficients to reduce the risk of overfitting. We considered three regularization methods: Ridge regression (L2 penalty, which shrinks coefficients toward zero but retains all features), Lasso regression (L1 penalty, which can shrink coefficients to exactly zero for automatic feature selection), and ElasticNet regression (which combines L1 and L2 penalties).

Model selection and evaluation were performed using nested cross-validation to prevent overfitting during hyperparameter selection^89^. In the outer loop, participants were partitioned into 10 folds (90% training, 10% testing). Within each outer fold, hyperparameters were optimized via an inner 10-fold cross-validation on the training data only, testing seven logarithmically spaced regularization strength parameters (*α*) from 0.001 to 1,000 (and five L1 values from 0.1 to 0.99 for ElasticNet). Model performance was evaluated on the held-out test data from the outer loop. Ridge regression demonstrated superior performance with an average cross-validated *R²* = .22 (**Figure 3C**).

For the final analysis, cerebellar parcel volumes were used to predict cerebral parcel volumes using Ridge regression with a global *α* = 100. This global *α* was determined via 10-fold cross-validation in a multioutput Ridge model predicting all cerebral parcels simultaneously and aligns with the distribution of parcel-wise optimal *α* values, where over 80% of cerebral parcels individually favored *α* = 100. Statistical significance of individual cerebellar-cerebral associations was assessed via permutation testing: the target variable was shuffled 10,000 times while maintaining the predictor structure, the model was refitted for each permutation, and a null distribution of regression coefficients was generated for each cerebellar-cerebral pair. The observed coefficients were then compared against this null distribution to derive one-sided *p*-values, which were subsequently corrected for multiple comparisons across all parcel pairs using FDR^50^ at *q* = .05.

### Association with behavioral outcomes

We used PLS analysis to associate participants’ normative model-derived *z*-scores for each cerebellar parcel with their behavioral scores in the HCP-D dataset (see **Behavioral tasks**). PLS is a multivariate technique that identifies orthogonal *latent variables*, pairs of weighted linear combinations of variables, which maximally covary^58^. Here, each latent variable consists of a set of parcel weights, a corresponding set of behavioral weights, and an associated singular value that quantifies the covariance between cerebellar structure and behavioral outcomes explained by that component. It is important to note that PLS does not establish causal links between cerebellar growth and behavioral outcomes, does not identify specific univariate parcel-behavior associations, and does not rule out the possibility of other relationships between cerebellar parcels and behaviors^60^.

Prior to the analysis, behaviors were standardized and residualized for age and sex to be consistent with cerebellar normative *z*-scores. We evaluated up to ten latent variables, with this maximum number determined by the smaller of the two matrix dimensions (i.e., behaviors). For each component, we computed the percentage of total covariance explained. Statistical significance of each component was assessed using permutation testing (10,000 iterations), in which the behavioral data were randomly shuffled while preserving the structure of the cerebellar matrix. This procedure yielded a null distribution of singular values under the assumption of no brain-behavior association. Since only the first latent variable was statistically significant, other latent variables were not analyzed further.

Statistical significance of the empirical correlation between normative parcel *z*-scores and behavioral outcomes was determined through permutation testing with 10,000 iterations, where behavioral scores were randomly shuffled to create a null distribution assuming no brain-behavior relationships. To assess stability of the PLS approach, we applied bootstrap resampling (10,000 iterations) and calculated 95% CIs around the loading weights of parcel *z*-scores and behaviors (**Supplementary** Figure 13). We computed bootstrap ratios (BSR) as the ratio of each loading to its bootstrap standard error across iterations. BSR values approximate *z*-scores, with |BSR| > 2.57 indicating significant contribution at *p* < .01 (two-sided)^59^. Importantly, BSR values also reflect the relative salience (contribution strength) of each variable to the brain-behavior relationship, allowing us to rank variables by their importance^59^ (**Supplementary Table 9**). Variables were considered significant if (1) 95% CI did not include zero, and (2) |BSR| > 2.57. Lastly, we employed 10-fold cross-validation on the PLS model. Specifically, the PLS model was fitted on the training set (90% of the data), and its performance was evaluated on the held-out test set (10% of the data). This procedure was repeated ten times with different random splits to assess the model’s generalizability across different subsets of participants. Correlations between cerebellar parcels and behavioral scores were separately calculated in training and test sets.

Finally, we assessed how cerebellar normative growth was associated with each behavior in relation to cerebral growth. Normative *z*-scores from the functional fusion cerebellar models and the DK cerebral models were used as multivariate predictors of individual behavioral measures. Cerebellar and cerebral parcels were analyzed separately and in combination by concatenating their respective features. For each behavioral outcome, we employed regularized regression to associate individual participants’ parcel *z*-scores (cerebellum-only, cerebral cortex-only, or combined cerebello-cerebral) with their behavioral scores, capturing distributed brain-behavior relationships. We compared three regularization methods: Ridge regression (L2 penalty), Lasso regression (L1 penalty), and ElasticNet regression (combined L1 and L2 penalties). Model selection and evaluation were performed using the same nested cross-validation procedure described in the previous section (**Association with normative trajectories of the cerebral cortex**). ElasticNet consistently achieved comparable or superior *R²* scores relative to Ridge and Lasso regression across all feature sets and behavioral outcomes (**Figure 4E**, left) and was therefore selected over other methods. Performance differences between cerebellar-only, cerebral cortex-only, and combined feature sets were evaluated using pairwise Wilcoxon signed-rank tests across the ten cross-validation folds.

### Data availability

The present study used existing developmental data from the Lifespan BCP (<u>nda.nih.gov/edit_collection.html?id=2848</u>) and the Lifespan 2.0 HCP-D release (nda.nih.gov/general-query.html?q=query=featured-datasets:HCP%20Aging%20and%20Development). Neuroimaging and behavioral data are publicly available for download from the provided links. The cerebellar functional atlases are available on GitHub (<u>github.com/DiedrichsenLab/cerebellar_atlases</u>). The cerebellar growth models for lobular and functional parcels are also available on GitHub (github.com/kmanoli/NormCerebellum).

### Code availability

Image preprocessing leveraged open-source software (iBEAT V2.0^82^: github.com/iBEAT-V2/iBEAT-V2.0-Docker; HCP minimal preprocessing pipeline^81^: github.com/Washington-University/HCPpipelines). Parcellations for the cerebellum and the cerebral cortex were also derived by openly available tools (ACAPULCO^43^: gitlab.com/shuohan/acapulco; FSL: fsl.fmrib.ox.ac.uk/fsl/docs/#/; FreeSurfer: surfer.nmr.mgh.harvard.edu). ITK-SNAP^84^ for manual correction of segmentations is freely available online (itksnap.org/pmwiki/pmwiki.php). Normative modeling code is available via the PCNtoolkit^25,44,45^ (github.com/amarquand/PCNtoolkit). Scripts for all processing and analysis pipelines used in this study can be found on GitHub (github.com/kmanoli/NormCerebellum).

## Supporting information

Supplementary information for manuscript.

## Acknowledgements

This work is supported by the Max Planck Society. AM was also funded by the German Academic Scholarship Foundation (Studienstiftung des deutschen Volkes) and the Lise Meitner Excellence Program (awarded to SLV). SLV and NM were also funded in part by Helmholtz Association’s Initiative and Networking Fund under the Helmholtz International Lab grant agreement InterLabs-0015, and the Canada First Research Excellence Fund (CFREF Competition 2, 2015-2016) awarded to the Healthy Brains, Healthy Lives initiative at McGill University, through the Helmholtz International BigBrain Analytics and Learning Laboratory (HIBALL). CGW was supported by an ERC Starting Grant (REPRESENT 101117806). JD and AFM were supported by the Raynor Cerebellum Project. SLV was furthermore supported by the Lise Meitner Excellence Program, the Jacobs Foundation Research Fellowship and the Hector Foundation Research Development Award.

## Author contributions

AM, NM, and SLV originally conceptualized and designed the study. AM curated, preprocessed, and segmented the data with support from NM, FH, TT, and LA. AM, NM, and LA performed image quality control under the supervision of SLV. AM, NM, and CY performed manual correction of cerebellar segmentations under the supervision of SLV. AM and NS set up and performed the normative modeling analyses. AM performed all additional analyses. AAAB and AFM provided expertise on normative modeling. JD provided expertise on functional parcellation of the cerebellum. SLV provided general project supervision. AM created the visualizations and wrote the original draft. All authors contributed to the interpretation of results. All authors edited and reviewed the final manuscript.

## References

1. Ito, M. Control of mental activities by internal models in the cerebellum. Nat. Rev. Neurosci. 9, 304–313 (2008).

2. Popa, L. S. & Ebner, T. J. Cerebellum, Predictions and Errors. Front. Cell. Neurosci. 12, (2019).

3. Tanaka, H., Ishikawa, T., Lee, J. & Kakei, S. The Cerebro-Cerebellum as a Locus of Forward Model: A Review. Front. Syst. Neurosci. 14, (2020).

4. King, M., Hernandez-Castillo, C. R., Poldrack, R. A., Ivry, R. B. & Diedrichsen, J. Functional boundaries in the human cerebellum revealed by a multi-domain task battery. Nat. Neurosci. 22, 1371–1378 (2019).

5. Manto, M. et al. Consensus Paper: Roles of the Cerebellum in Motor Control—The Diversity of Ideas on Cerebellar Involvement in Movement. Cerebellum Lond. Engl. 11, 457–487 (2012).

6. Nettekoven, C. et al. A hierarchical atlas of the human cerebellum for functional precision mapping. Nat. Commun. 15, 8376 (2024).

7. Van Overwalle, F. et al. Consensus Paper: Cerebellum and Social Cognition. The Cerebellum 19, 833–868 (2020).

8. Krienen, F. M. & Buckner, R. L. Segregated fronto-cerebellar circuits revealed by intrinsic functional connectivity. Cereb. Cortex N. Y. N 1991 19, 2485–2497 (2009).

9. Palesi, F. et al. Contralateral cerebello-thalamo-cortical pathways with prominent involvement of associative areas in humans in vivo. Brain Struct. Funct. 220, 3369–3384 (2015).

10. Palesi, F. et al. Contralateral cortico-ponto-cerebellar pathways reconstruction in humans in vivo: implications for reciprocal cerebro-cerebellar structural connectivity in motor and non-motor areas. Sci. Rep. 7, 12841 (2017).

11. Sokolov, A. A., Erb, M., Grodd, W. & Pavlova, M. A. Structural Loop Between the Cerebellum and the Superior Temporal Sulcus: Evidence from Diffusion Tensor Imaging. Cereb. Cortex 24, 626–632 (2014).

12. Palacios, E. R., Chadderton, P., Friston, K. & Houghton, C. Cerebellar state estimation enables resilient coupling across behavioural domains. Sci. Rep. 14, 6641 (2024).

13. Van Overwalle, F., Manto, M., Leggio, M. & Delgado-García, J. M. The sequencing process generated by the cerebellum crucially contributes to social interactions. Med. Hypotheses 128, 33–42 (2019).

14. Limperopoulos, C. et al. Late Gestation Cerebellar Growth Is Rapid and Impeded by Premature Birth. Pediatrics 115, 688–695 (2005).

15. ten Donkelaar, H. J., Lammens, M., Wesseling, P., Thijssen, H. O. M. & Renier, W. O. Development and developmental disorders of the human cerebellum. J. Neurol. 250, 1025– 1036 (2003).

16. Wang, V. Y. & Zoghbi, H. Y. Genetic regulation of cerebellar development. Nat. Rev. Neurosci. 2, 484–491 (2001).

17. King, M. The big role of the ‘little brain’: exploring the developing cerebellum and its role in cognition. Curr. Opin. Behav. Sci. 54, 101301 (2023).

18. Olson, I. R., Hoffman, L. J., Jobson, K. R., Popal, H. S. & Wang, Y. Little brain, little minds: The big role of the cerebellum in social development. Dev. Cogn. Neurosci. 60, 101238 (2023).

19. Moberget, T. et al. Cerebellar volume and cerebellocerebral structural covariance in schizophrenia: a multisite mega-analysis of 983 patients and 1349 healthy controls. Mol. Psychiatry 23, 1512–1520 (2018).

20. Stoodley, C. J. The Cerebellum and Neurodevelopmental Disorders. The Cerebellum 15, 34– 37 (2016).

21. Wang, S. S.-H., Kloth, A. D. & Badura, A. The cerebellum, sensitive periods, and autism. Neuron 83, 518–532 (2014).

22. Reumers, S. F. I. et al. Cognition in cerebellar disorders: What’s in the profile? A systematic review and meta-analysis. J. Neurol. 272, 250 (2025).

23. Stoodley, C. J. & Limperopoulos, C. Structure–function relationships in the developing cerebellum: evidence from early-life cerebellar injury and neurodevelopmental disorders. Semin. Fetal. Neonatal Med. 21, 356–364 (2016).

24. Bethlehem, R. a. I., et al. Brain charts for the human lifespan. Nature 604, 525–533 (2022).

25. Rutherford, S. et al. The normative modeling framework for computational psychiatry. Nat. Protoc. 17, 1711–1734 (2022).

26. Kia, S. M. et al. Closing the life-cycle of normative modeling using federated hierarchical Bayesian regression. PLOS ONE 17, e0278776 (2022).

27. Rutherford, S. et al. Charting brain growth and aging at high spatial precision. eLife 11, e72904 (2022).

28. Gaiser, C. et al. Population-wide cerebellar growth models of children and adolescents. Nat. Commun. 15, 2351 (2024).

29. Kim, M. et al. Mapping cerebellar anatomical heterogeneity in mental and neurological illnesses. Nat. Ment. Health 2, 1196–1207 (2024).

30. Wang, Y. et al. Longitudinal development of the cerebellum in human infants during the first 800 days. Cell Rep. 42, 112281 (2023).

31. Constantino JN, & Gruber CP. (2012) Social Responsiveness Scale–Second Edition (SRS-2). Torrance, CA: Western Psychological Services.

32. Manoli, A., Van Overwalle, F., Grosse Wiesmann, C. & Valk, S. L. Functional recruitment and connectivity of the cerebellum is associated with the emergence of Theory of Mind in early childhood. Nat. Commun. 16, 5273 (2025).

33. Uquillas, F. d’Oleire et al. The Cerebellum Plays a Protective Role in Cognitive Aging and Disease: Insights from a Multi-Cohort Study. Alzheimers Dement. 20, e085743 (2025).

34. Wang, D., Buckner, R. L. & Liu, H. Cerebellar asymmetry and its relation to cerebral asymmetry estimated by intrinsic functional connectivity. J. Neurophysiol. 109, 46–57 (2013).

35. Wang, Z. et al. Structural covariation between cerebellum and neocortex intrinsic structural covariation links cerebellum subregions to the cerebral cortex. J. Neurophysiol. 132, 849–869 (2024).

36. Buckner, R. L., Krienen, F. M., Castellanos, A., Diaz, J. C. & Yeo, B. T. T. The organization of the human cerebellum estimated by intrinsic functional connectivity. J. Neurophysiol. 106, 2322–2345 (2011).

37. Guell, X., Schmahmann, J. D., Gabrieli, J. D. & Ghosh, S. S. Functional gradients of the cerebellum. eLife 7, e36652 (2018).

38. Marek, S. et al. Spatial and Temporal Organization of the Individual Human Cerebellum. Neuron 100, 977–993.e7 (2018).

39. Magielse, N. et al. A Bias-Accounting Meta-Analytic Approach Refines and Expands the Cerebellar Behavioral Topography. 2024.10.31.621398 Preprint at 10.1101/2024.10.31.621398 (2025).

40. Stoodley, C. J. & Schmahmann, J. D. Functional topography in the human cerebellum: a meta-analysis of neuroimaging studies. NeuroImage 44, 489–501 (2009).

41. Howell, B. R. et al. The UNC/UMN Baby Connectome Project (BCP): An Overview of the Study Design and Protocol Development. NeuroImage 185, 891–905 (2019).

42. Somerville, L. H. et al. The Lifespan Human Connectome Project in Development: A large-scale study of brain connectivity development in 5–21 year olds. NeuroImage 183, 456–468 (2018).

43. Han, S., Carass, A., He, Y. & Prince, J. L. Automatic cerebellum anatomical parcellation using U-Net with locally constrained optimization. NeuroImage 218, 116819 (2020).

44. de Boer, A. A. A. et al. Non-Gaussian normative modelling with hierarchical Bayesian regression. Imaging Neurosci. 2, 1–36 (2024).

45. Marquand, A. F., Rezek, I., Buitelaar, J. & Beckmann, C. F. Understanding Heterogeneity in Clinical Cohorts Using Normative Models: Beyond Case-Control Studies. Biol. Psychiatry 80, 552–561 (2016).

46. Brooks, S. P. & Gelman, A. General Methods for Monitoring Convergence of Iterative Simulations. J. Comput. Graph. Stat. 7, 434–455 (1998).

47. Sun, L. et al. Human lifespan changes in the brain’s functional connectome. Nat. Neurosci. 28, 891–901 (2025).

48. Yates, T. S., Ellis, C. T. & Turk-Browne, N. B. Emergence and organization of adult brain function throughout child development. NeuroImage 226, 117606 (2021).

49. Desikan, R. S. et al. An automated labeling system for subdividing the human cerebral cortex on MRI scans into gyral based regions of interest. NeuroImage 31, 968–980 (2006).

50. Benjamini, Y. & Hochberg, Y. Controlling the False Discovery Rate: A Practical and Powerful Approach to Multiple Testing. J. R. Stat. Soc. Ser. B Methodol. 57, 289–300 (1995).

51. Utevsky, A. V., Smith, D. V. & Huettel, S. A. Precuneus Is a Functional Core of the Default-Mode Network. J. Neurosci. 34, 932–940 (2014).

52. Fransson, P. & Marrelec, G. The precuneus/posterior cingulate cortex plays a pivotal role in the default mode network: Evidence from a partial correlation network analysis. NeuroImage 42, 1178–1184 (2008).

53. Takehara-Nishiuchi, K. Entorhinal cortex and consolidated memory. Neurosci. Res. 84, 27– 33 (2014).

54. Badre, D. Cognitive control, hierarchy, and the rostro–caudal organization of the frontal lobes. Trends Cogn. Sci. 12, 193–200 (2008).

55. Deschamps, I., Baum, S. R. & Gracco, V. L. On the role of the supramarginal gyrus in phonological processing and verbal working memory: Evidence from rTMS studies. Neuropsychologia 53, 39–46 (2014).

56. Spasojević, G., Malobabic, S., Pilipović-Spasojević, O., Djukić-Macut, N. & Maliković, A. Morphology and digitally aided morphometry of the human paracentral lobule. Folia Morphol. 72, 10–16 (2013).

57. Nelson, A. J. & Chen, R. Digit somatotopy within cortical areas of the postcentral gyrus in humans. Cereb. Cortex N. Y. N 1991 18, 2341–2351 (2008).

58. Krishnan, A., Williams, L. J., McIntosh, A. R. & Abdi, H. Partial Least Squares (PLS) methods for neuroimaging: A tutorial and review. NeuroImage 56, 455–475 (2011).

59. McIntosh, A. R. & Lobaugh, N. J. Partial least squares analysis of neuroimaging data: applications and advances. NeuroImage 23, S250–S263 (2004).

60. Hansen, J. Y. et al. Mapping neurotransmitter systems to the structural and functional organization of the human neocortex. Nat. Neurosci. 25, 1569–1581 (2022).

61. Tiemeier, H. et al. Cerebellum development during childhood and adolescence: A longitudinal morphometric MRI study. NeuroImage 49, 63–70 (2010).

62. Brossard-Racine, M. & Limperopoulos, C. Normal Cerebellar Development by Qualitative and Quantitative MR Imaging: From the Fetus to the Adolescent. Neuroimaging Clin. N. Am. 26, 331–339 (2016).

63. Balsters, J. H. et al. Evolution of the cerebellar cortex: The selective expansion of prefrontal-projecting cerebellar lobules. NeuroImage 49, 2045–2052 (2010).

64. Magielse, N. et al. Phylogenetic comparative analysis of the cerebello-cerebral system in 34 species highlights primate-general expansion of cerebellar crura I-II. Commun. Biol. 6, 1188 (2023).

65. Liu, X., d’Oleire Uquillas, F., Viaene, A. N., Zhen, Z. & Gomez, J. A multifaceted gradient in human cerebellum of structural and functional development. Nat. Neurosci. 25, 1129–1133 (2022).

66. Klein, A. P., Ulmer, J. L., Quinet, S. A., Mathews, V. & Mark, L. P. Nonmotor Functions of the Cerebellum: An Introduction. AJNR Am. J. Neuroradiol. 37, 1005–1009 (2016).

67. Sathyanesan, A. et al. Emerging connections between cerebellar development, behaviour and complex brain disorders. Nat. Rev. Neurosci. 20, 298–313 (2019).

68. d’Oleire Uquillas, F., et al. Multimodal evidence for cerebellar influence on cortical development in autism: structural growth amidst functional disruption. Mol. Psychiatry 30, 1558–1572 (2025).

69. Limperopoulos, C. et al. Injury to the Premature Cerebellum: Outcome is Related to Remote Cortical Development. Cereb. Cortex 24, 728–736 (2014).

70. Mariën, P. et al. Consensus paper: Language and the cerebellum: an ongoing enigma. Cerebellum Lond. Engl. 13, 386–410 (2014).

71. Jobson, K. R. et al. Language and the Cerebellum: Structural Connectivity to the Eloquent Brain. Neurobiol. Lang. Camb. Mass 5, 652–675 (2024).

72. Stoodley, C. J. & Tsai, P. T. Adaptive Prediction for Social Contexts: The Cerebellar Contribution to Typical and Atypical Social Behaviors. Annu. Rev. Neurosci. 44, 475–493 (2021).

73. Van Overwalle, F. et al. The Involvement of the Posterior Cerebellum in Reconstructing and Predicting Social Action Sequences. The Cerebellum 21, 733–741 (2022).

74. Fatemi, S. H. et al. Consensus Paper: Pathological Role of the Cerebellum in Autism. The Cerebellum 11, 777–807 (2012).

75. Rogers, T. D. et al. Is autism a disease of the cerebellum? An integration of clinical and pre-clinical research. Front. Syst. Neurosci. 7, (2013).

76. Diedrichsen, J. A spatially unbiased atlas template of the human cerebellum. NeuroImage 33, 127–138 (2006).

77. Diedrichsen, J. & Zotow, E. Surface-Based Display of Volume-Averaged Cerebellar Imaging Data. PLOS ONE 10, e0133402 (2015).

78. Lyu, W. et al. Functional development of the human cerebellum from birth to age five. Nat. Commun. 16, 6350 (2025).

79. Pfefferbaum, A. & Sullivan, E. V. Cross-sectional versus longitudinal estimates of age-related changes in the adult brain: overlaps and discrepancies. Neurobiol. Aging 36, 2563–2567 (2015).

80. Harms, M. P. et al. Extending the Human Connectome Project across ages: Imaging protocols for the Lifespan Development and Aging projects. NeuroImage 183, 972–984 (2018).

81. Glasser, M. F. et al. The minimal preprocessing pipelines for the Human Connectome Project. NeuroImage 80, 105–124 (2013).

82. Wang, L. et al. iBEAT V2.0: a multisite-applicable, deep learning-based pipeline for infant cerebral cortical surface reconstruction. Nat. Protoc. 18, 1488–1509 (2023).

83. Fonov, V., Evans, A., McKinstry, R., Almli, C. & Collins, D. Unbiased nonlinear average age-appropriate brain templates from birth to adulthood. NeuroImage 47, S102 (2009).

84. Yushkevich, P. A. et al. User-guided 3D active contour segmentation of anatomical structures: significantly improved efficiency and reliability. NeuroImage 31, 1116–1128 (2006).

85. Fischl, B., Liu, A. & Dale, A. M. Automated manifold surgery: constructing geometrically accurate and topologically correct models of the human cerebral cortex. IEEE Trans. Med. Imaging 20, 70–80 (2001).

86. Rosen, A. F. G. et al. Quantitative Assessment of Structural Image Quality. NeuroImage 169, 407–418 (2018).

87. Remiszewski, N. et al. Contrasting Case-Control and Normative Reference Approaches to Capture Clinically Relevant Structural Brain Abnormalities in Patients With First-Episode Psychosis Who Are Antipsychotic Naive. JAMA Psychiatry 79, 1133–1138 (2022).

88. Zöllei, L., Iglesias, J. E., Ou, Y., Grant, P. E. & Fischl, B. Infant FreeSurfer: An automated segmentation and surface extraction pipeline for T1-weighted neuroimaging data of infants 0– 2 years. NeuroImage 218, 116946 (2020).

89. Varma, S. & Simon, R. Bias in error estimation when using cross-validation for model selection. BMC Bioinformatics 7, 91 (2006).

