## Supplementary information for manuscript. for "Cerebellar growth is associated with domain-specific cerebral maturation and socio-linguistic behavioral outcomes"

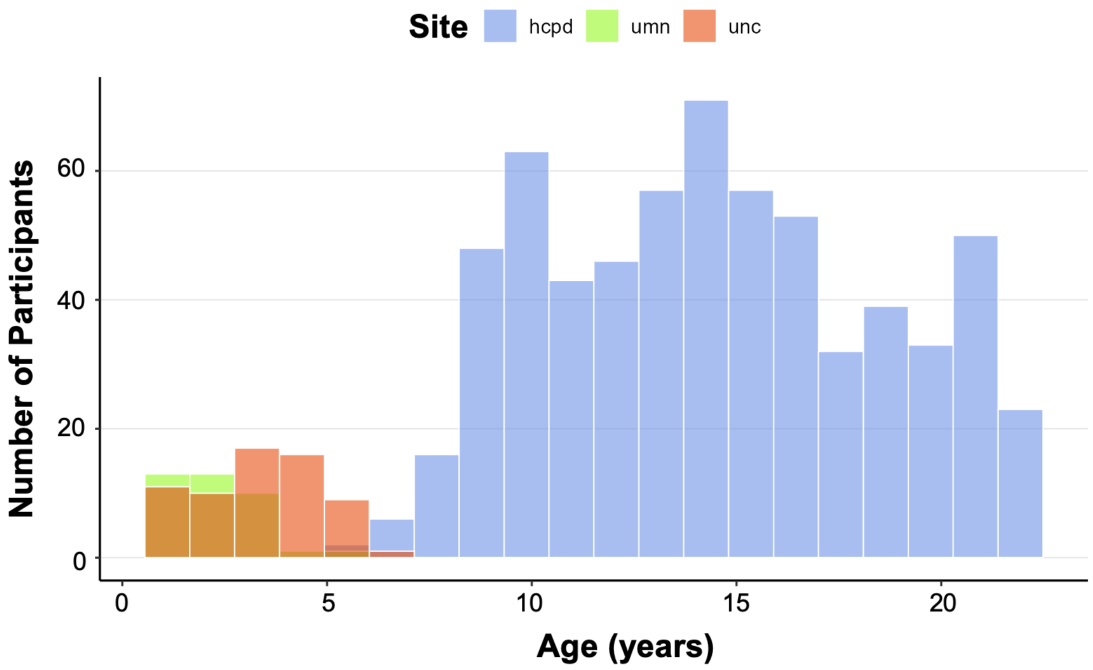

**Supplementary Figure 1.** Age distribution per scanner site. Abbreviations: hcpd = Human Connectome Project Development; umn = Baby Connectome Project University of Minnesota sample; unc = Baby Connectome Project University of North Carolina sample.

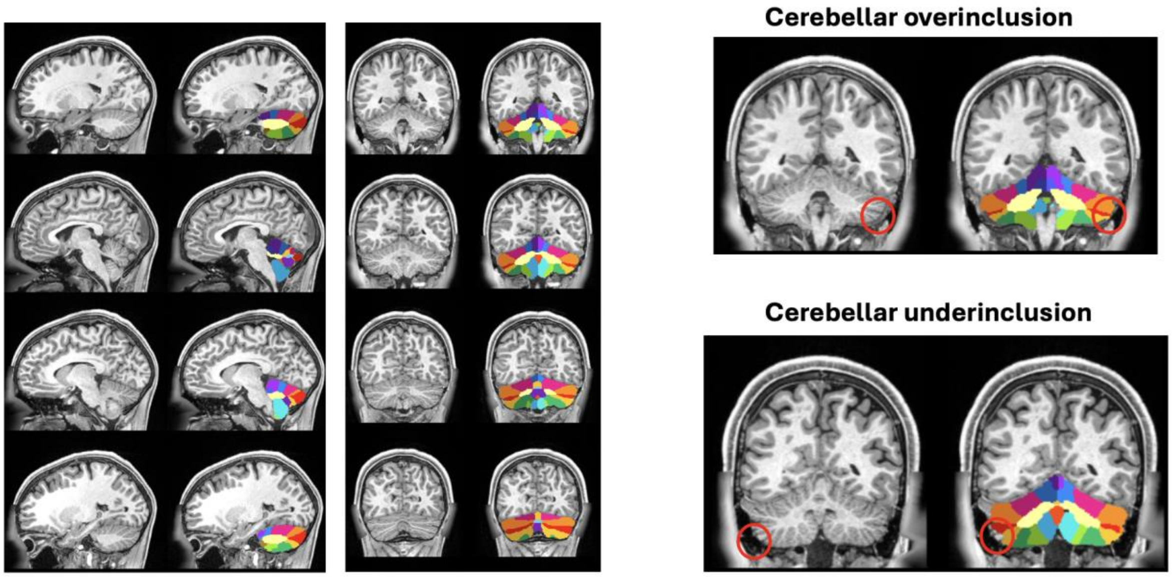

**Supplementary Figure 2.** Example lobular segmentations performed with the ACAPULCO algorithm. Left: Example ACAPULCO segmentation, with a native space lobular mask overlaid on a participant's T1-weighted image (sagittal and coronal views). Right: Example of cerebellar overinclusion (top; circled in red) and underinclusion (bottom; circled in red). For both types of segmentation errors, slice-wise manual correction of the mask was performed by tracing cerebellar fissures and folia (see **Methods: Cerebellar parcellation**). In the case of overinclusion, the ACAPULCO mask included parts of the cerebral cortex, non-brain infratentorial tissue, and/or lower skull and neck tissue. In the case of underinclusion, the ACAPULCO mask often missed parts of the cerebellar grey matter. Both errors were most common around postero-lateral cerebellar regions.

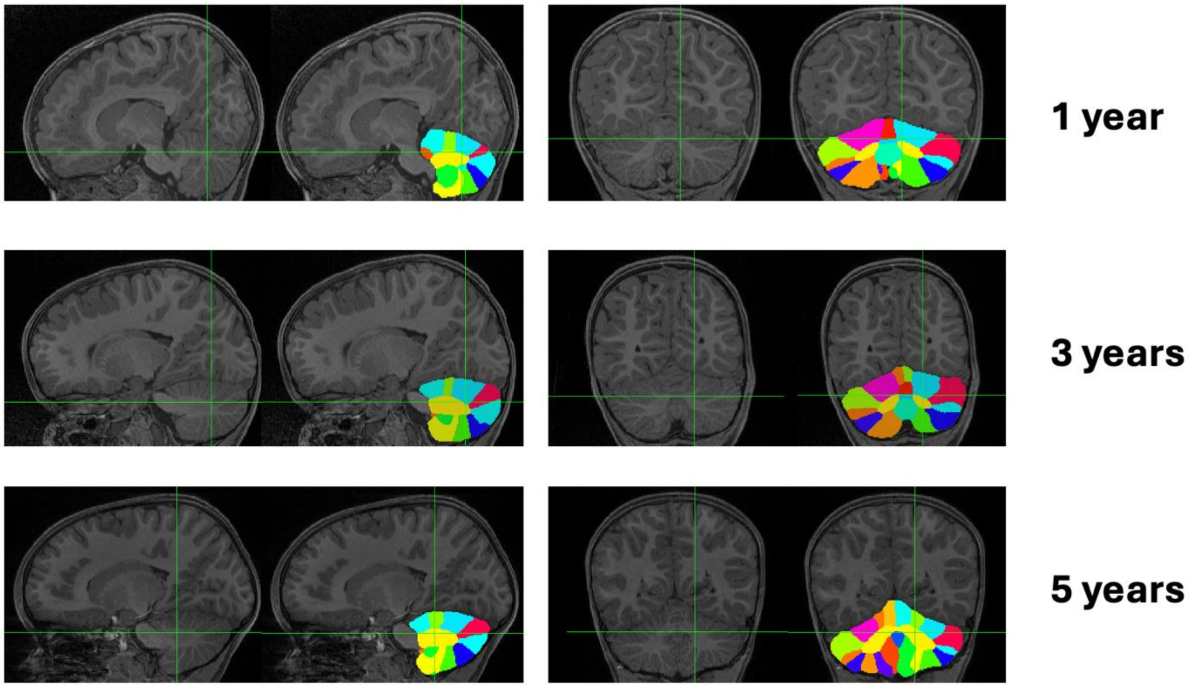

**Supplementary Figure 3.** Example lobular segmentations performed with the ACAPULCO algorithm in the BCP dataset (see **Methods: Cerebellar parcellation**). Native space ACAPULCO segmentation masks are overlaid on participants' T1-weighted images (sagittal and coronal views) in example timepoints (1, 3, and 5 years) across the BCP age range (1-5 years).

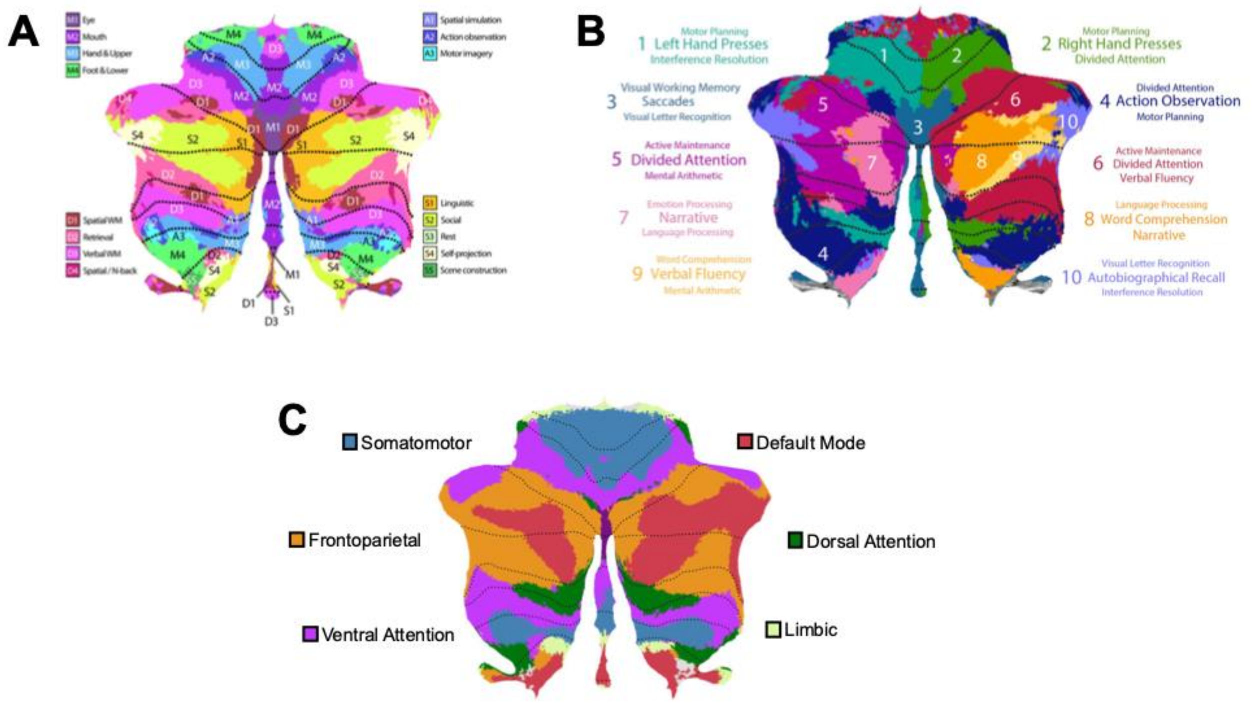

**Supplementary Figure 4.** Functional cerebellar parcellations. **A.** Functional fusion atlas, adapted with permission from Nettekoven et al. (2024). **B.** MDTB atlas, adapted with permission from King et al. (2019). **C.** Resting-state atlas, based on Buckner et al. (2011) and plotted via the openly available cerebellar atlas viewer (diedrichsenlab.org/imaging/AtlasViewer/index.htm).

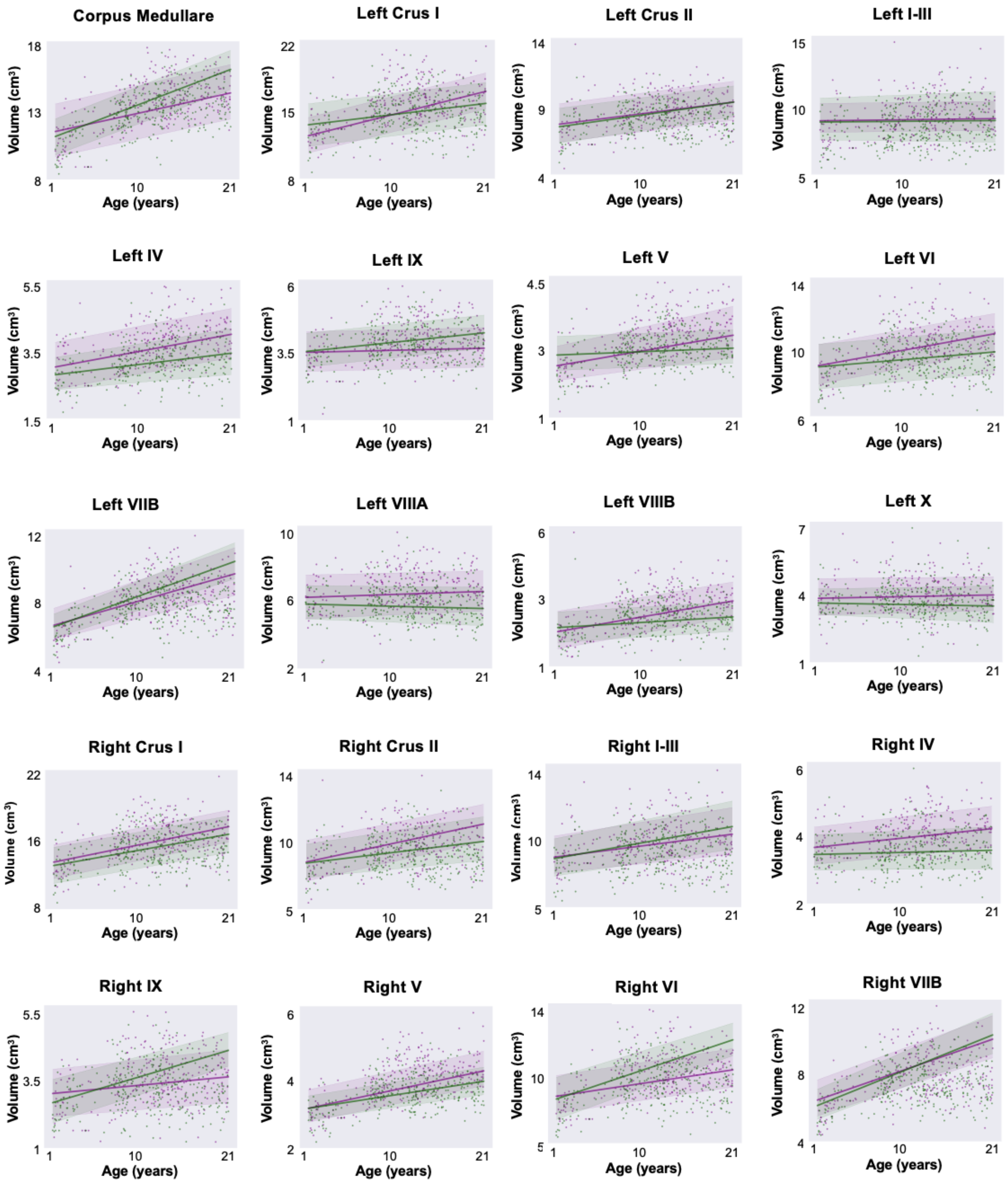

Corpus Medullare
Volume (cm³)
18
13
8
1
10
21
Age (years)
Left Crus I
22
15
8
Left Crus II
14
9
4
Left I-III
15
10
5
Left IV
5.5
3.5
1.5
Left IX
6
3.5
1
Left V
4.5
3
1
Left VI
14
10
6
Left VIIB
12
8
4
Left VIIIA
10
6
2
Left VIIIB
6
3
1
Left X
7
4
1
Right Crus I
22
16
8
Right Crus II
14
10
5
Right I-III
14
10
5
Right IV
6
4
2
Right IX
5.5
3.5
1
Right V
6
4
2
Right VI
14
10
5
Right VIIB
12
8
4

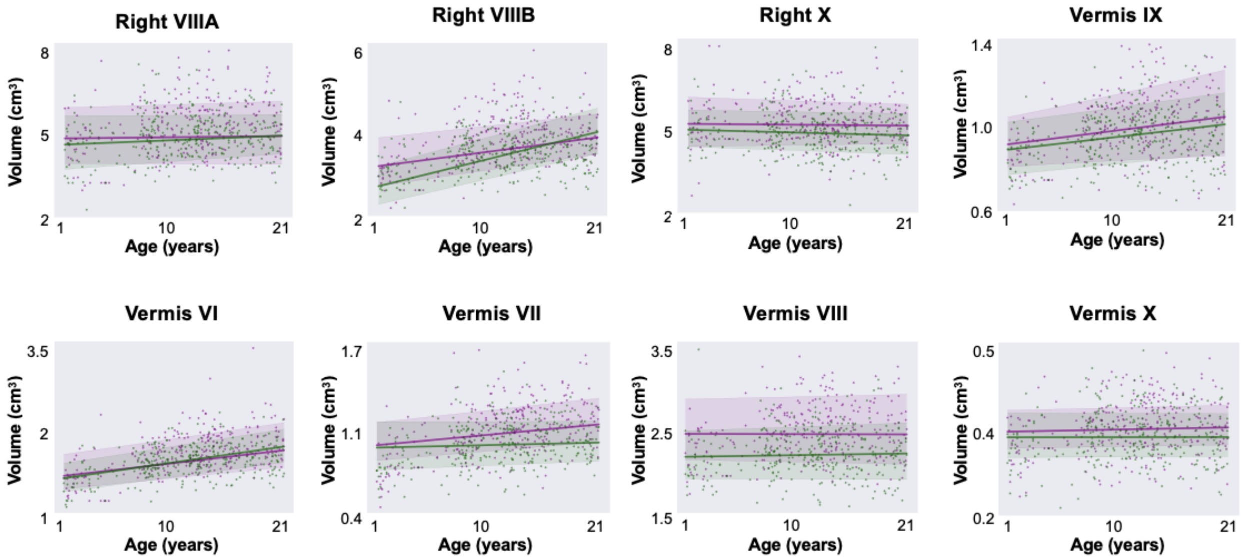

**Supplementary Figure 5.** Sex-stratified normative trajectories for all lobules in the ACAPULCO segmentation. Bold lines represent the mean trajectory per sex. Shaded areas represent the 95.5% confidence interval. Purple: male; green: female.

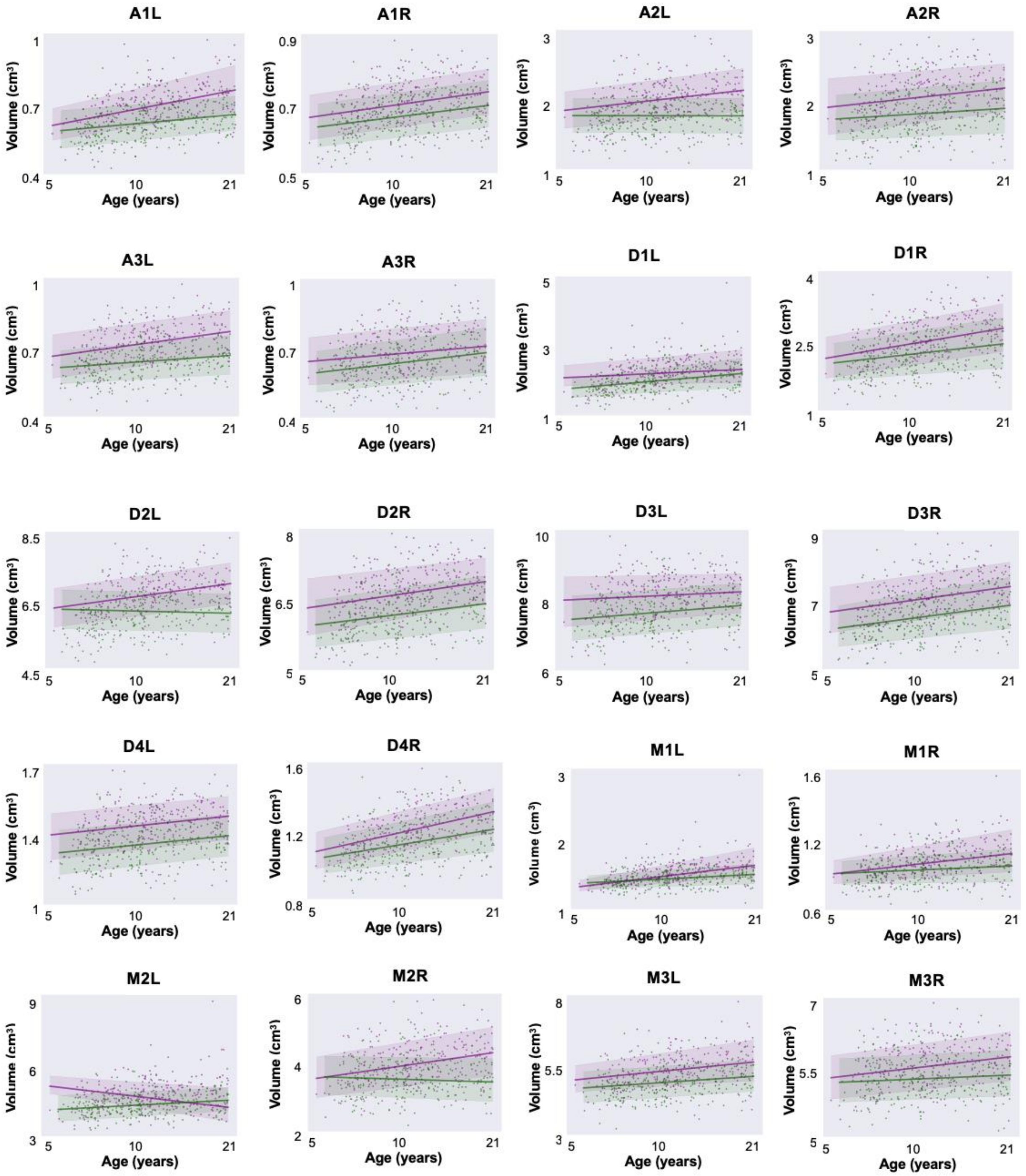
A1L
A1R
A2L
A2R
A3L
A3R
D1L
D1R
D2L
D2R
D3L
D3R
D4L
D4R
M1L
M1R
M2L
M2R
M3L
M3R
Volume (cm³)
Age (years)

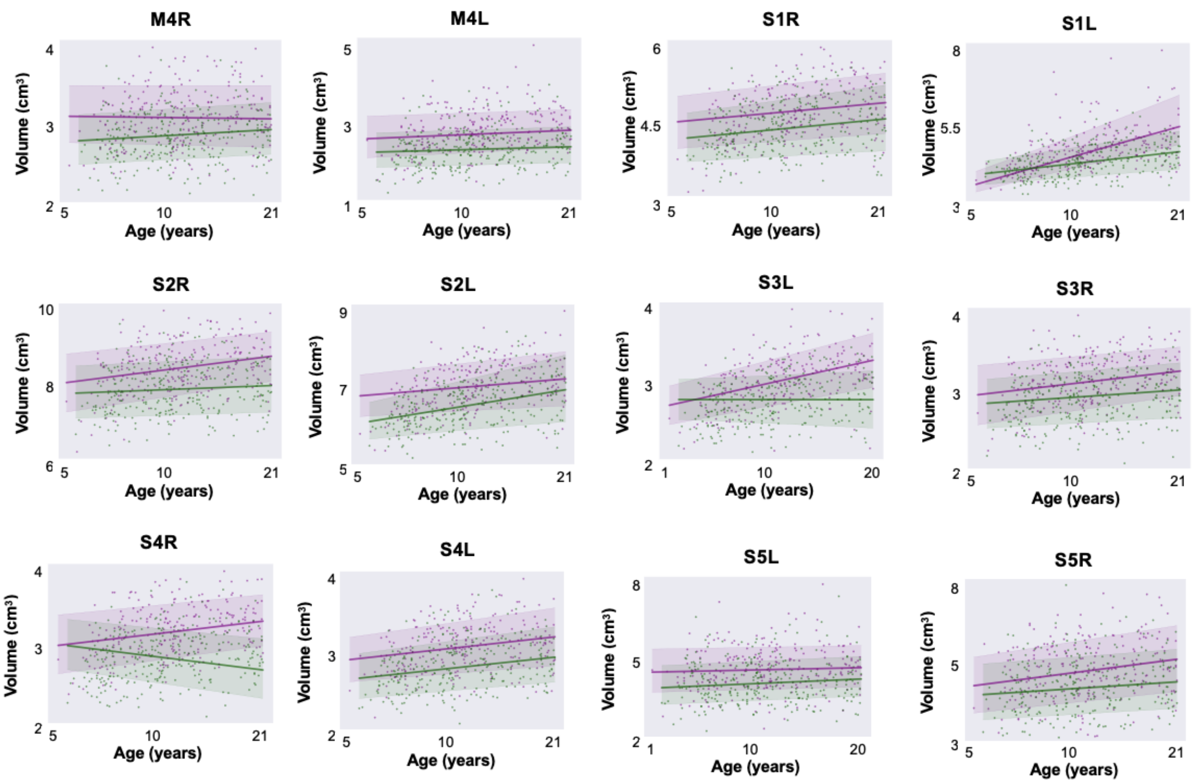

**Supplementary Figure 6.** Sex-stratified normative trajectories for all parcels in the functional fusion atlas. Bold lines represent the mean trajectory per sex. Shaded areas represent the 95.5% confidence interval. Purple: male; Green: female. Abbreviations: A = Action; D = Demand; M = Motor; S = Socio-Linguistic; L = left; R = right.

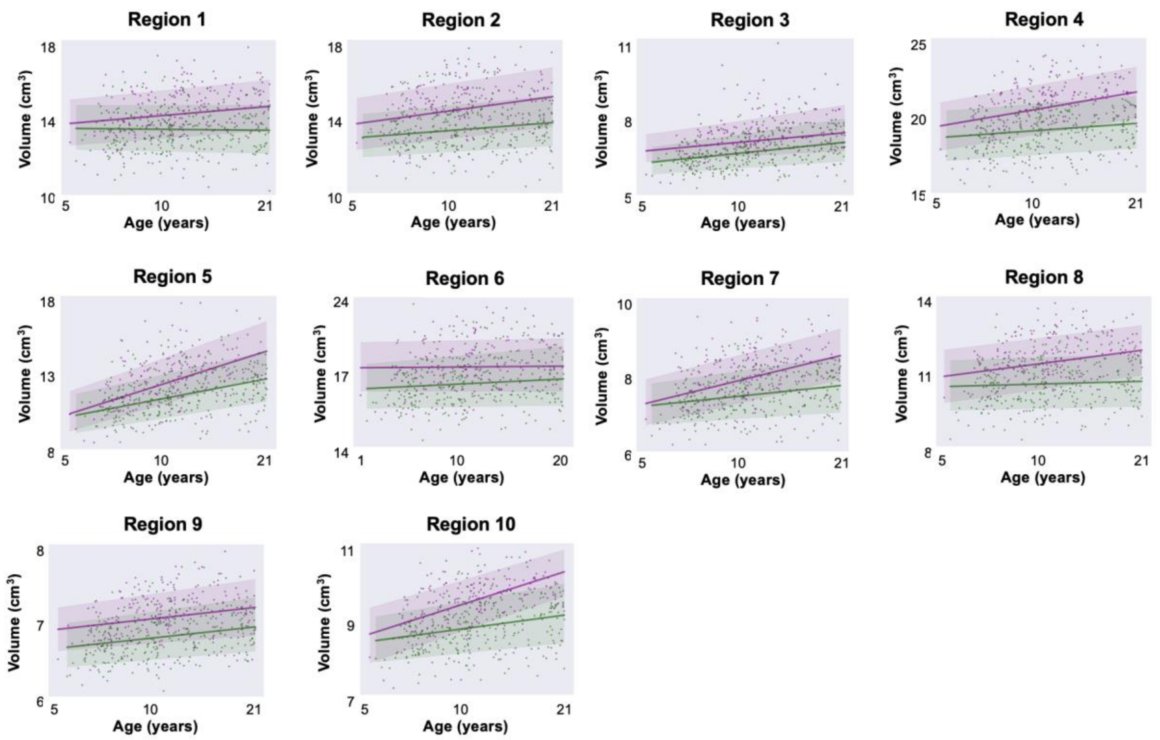

**Supplementary Figure 7.** Sex-stratified normative trajectories for all parcels in the MDTB atlas. Bold lines represent the mean trajectory per sex. Shaded areas represent the 95.5% confidence interval. Purple: male; green: female.

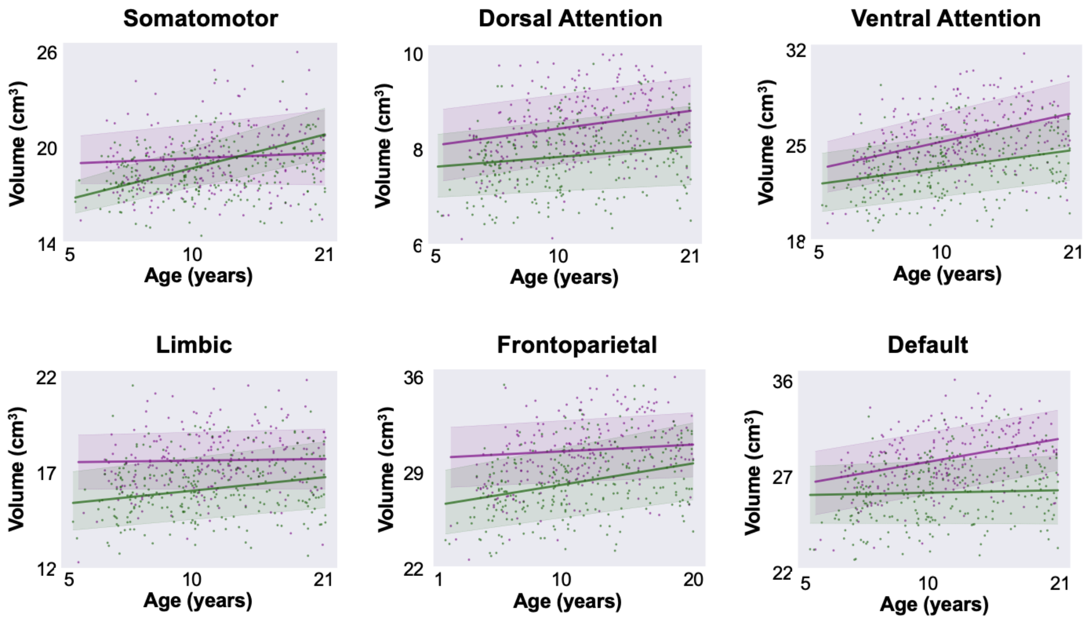

**Supplementary Figure 8.** Sex-stratified normative trajectories for all parcels in the resting-state atlas. Bold lines represent the mean trajectory per sex. Shaded areas represent the 95.5% confidence interval. Purple: male; green: female.

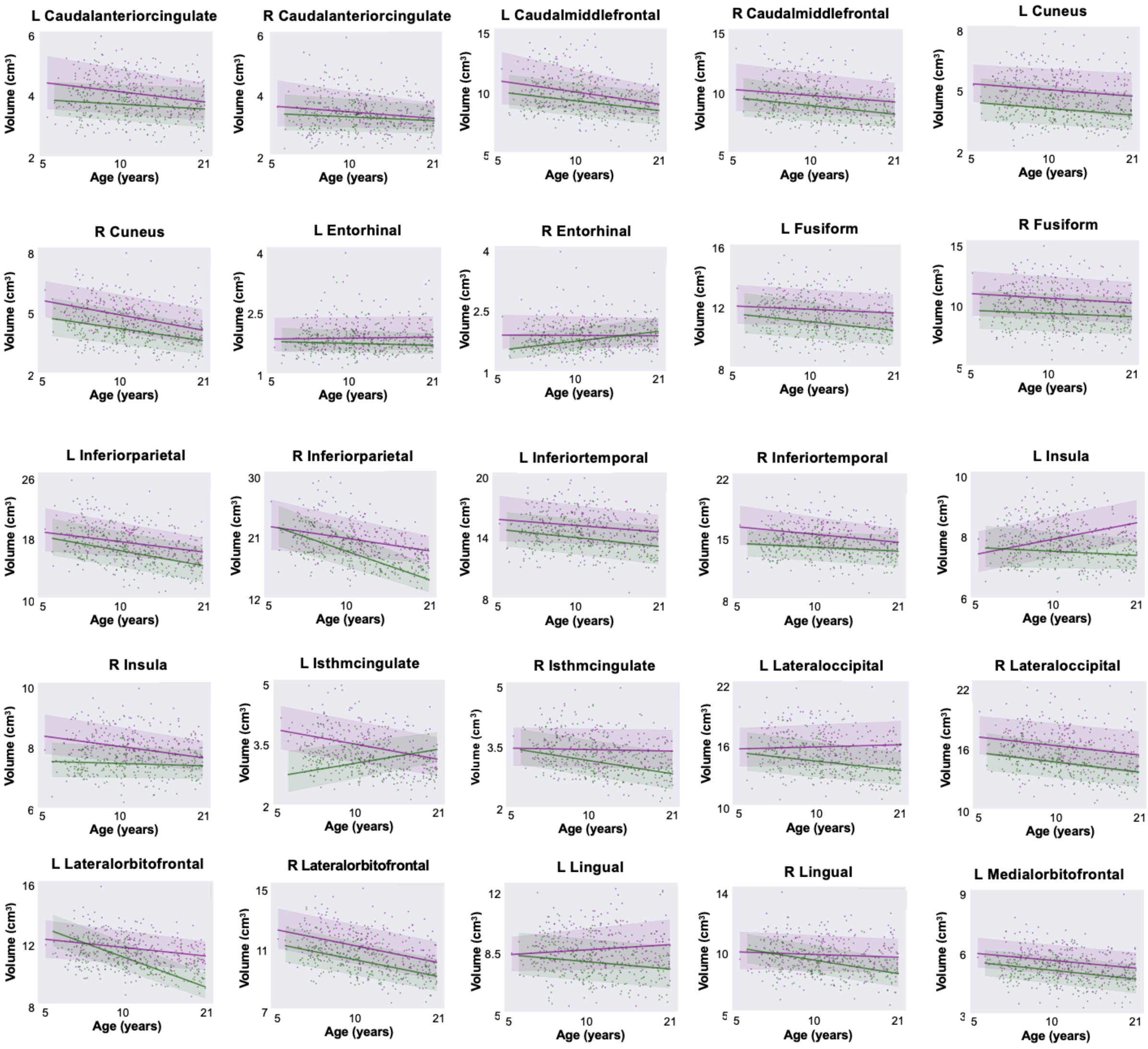
L Caudalanteriorcingulate
R Caudalanteriorcingulate
L Caudalmiddlefrontal
R Caudalmiddlefrontal
L Cuneus
R Cuneus
L Entorhinal
R Entorhinal
L Fusiform
R Fusiform
L Inferiorparietal
R Inferiorparietal
L Inferiortemporal
R Inferiortemporal
L Insula
R Insula
L Isthmcingulate
R Isthmcingulate
L Lateraloccipital
R Lateraloccipital
L Lateralorbitofrontal
R Lateralorbitofrontal
L Lingual
R Lingual
L Medialorbitofrontal
Volume (cm³)
Age (years)

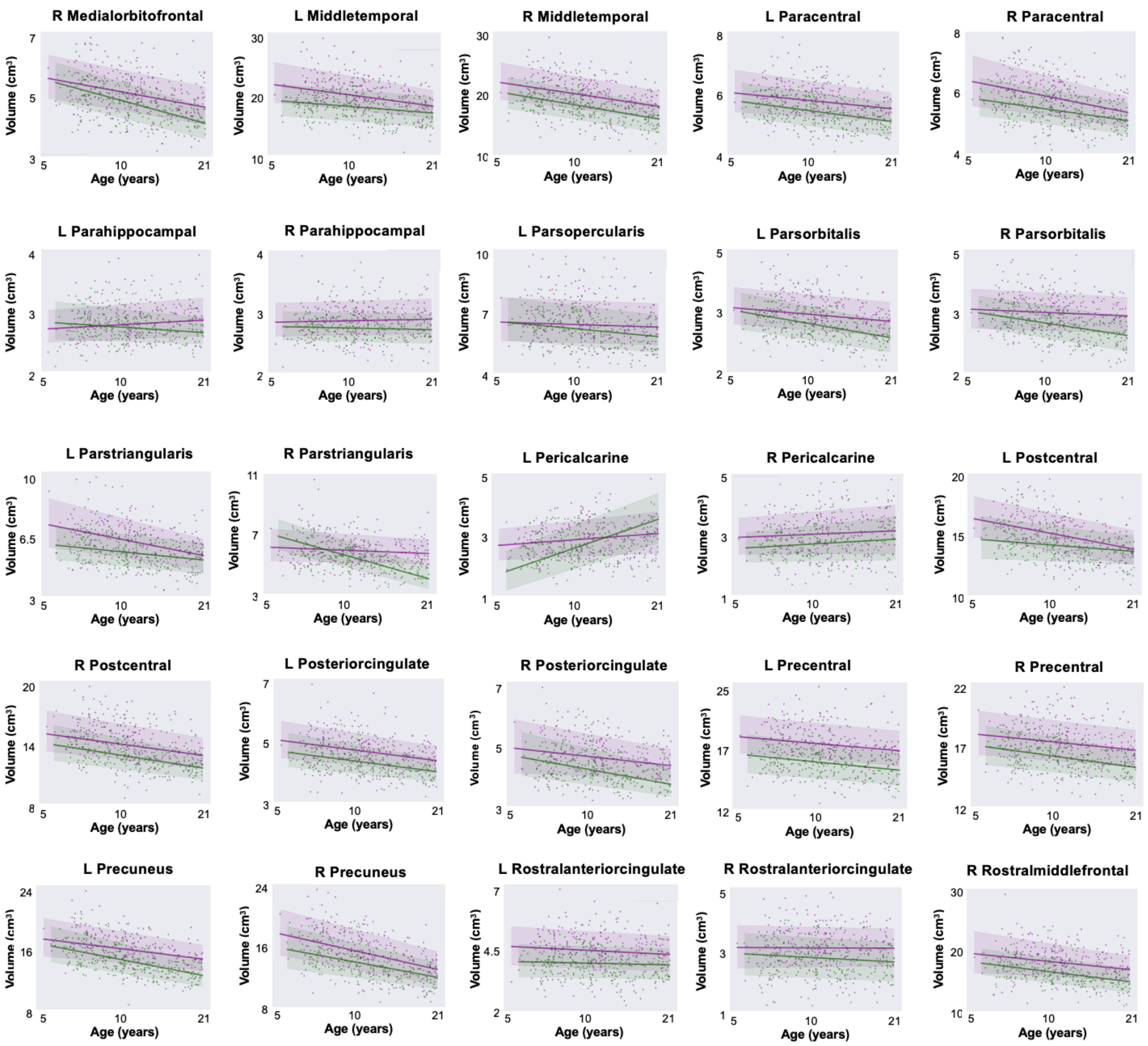
R Medialorbitofrontal
L Middletemporal
R Middletemporal
L Paracentral
R Paracentral
L Parahippocampal
R Parahippocampal
L Parsopercularis
L Parsorbitalis
R Parsorbitalis
L Parstriangularis
R Parstriangularis
L Pericalcarine
R Pericalcarine
L Postcentral
R Postcentral
L Posteriorcingulate
R Posteriorcingulate
L Precentral
R Precentral
L Precuneus
R Precuneus
L Rostralanteriorcingulate
R Rostralanteriorcingulate
R Rostralmiddlefrontal
Volume ($cm^3$)
Age (years)

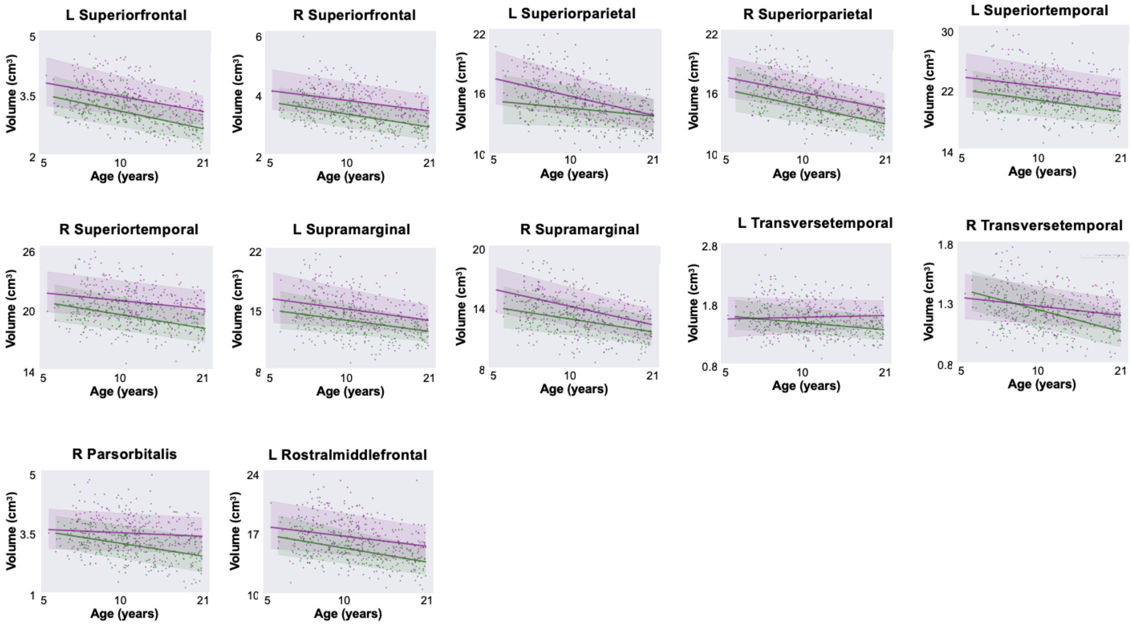

**Supplementary Figure 9.** Sex-stratified normative trajectories for all parcels in the DK atlas. Bold lines represent the mean trajectory per sex. Shaded areas represent the 95.5% confidence interval. Purple: male; green: female. Abbreviations: L = left; R = right.

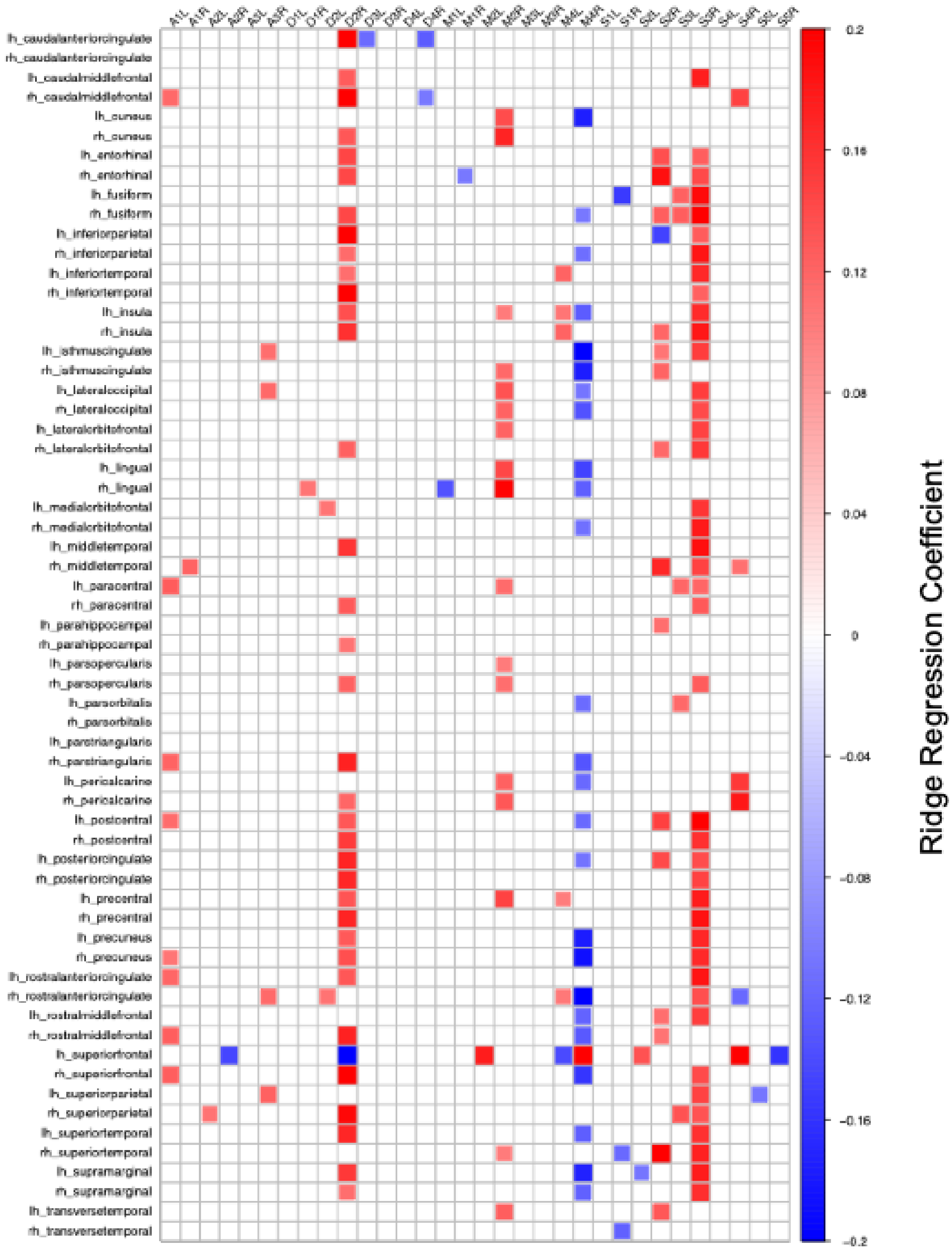

**Supplementary Figure 10.** Association of cerebellar (fusion atlas) and cerebral (DK) growth trajectories per DK hemisphere. The heatmap shows significant cerebro-cerebellar associations (10,000 permutations, false discovery rate (FDR)-corrected at $q$ = .05). Abbreviations: lh = left hemisphere; rh = right hemisphere.

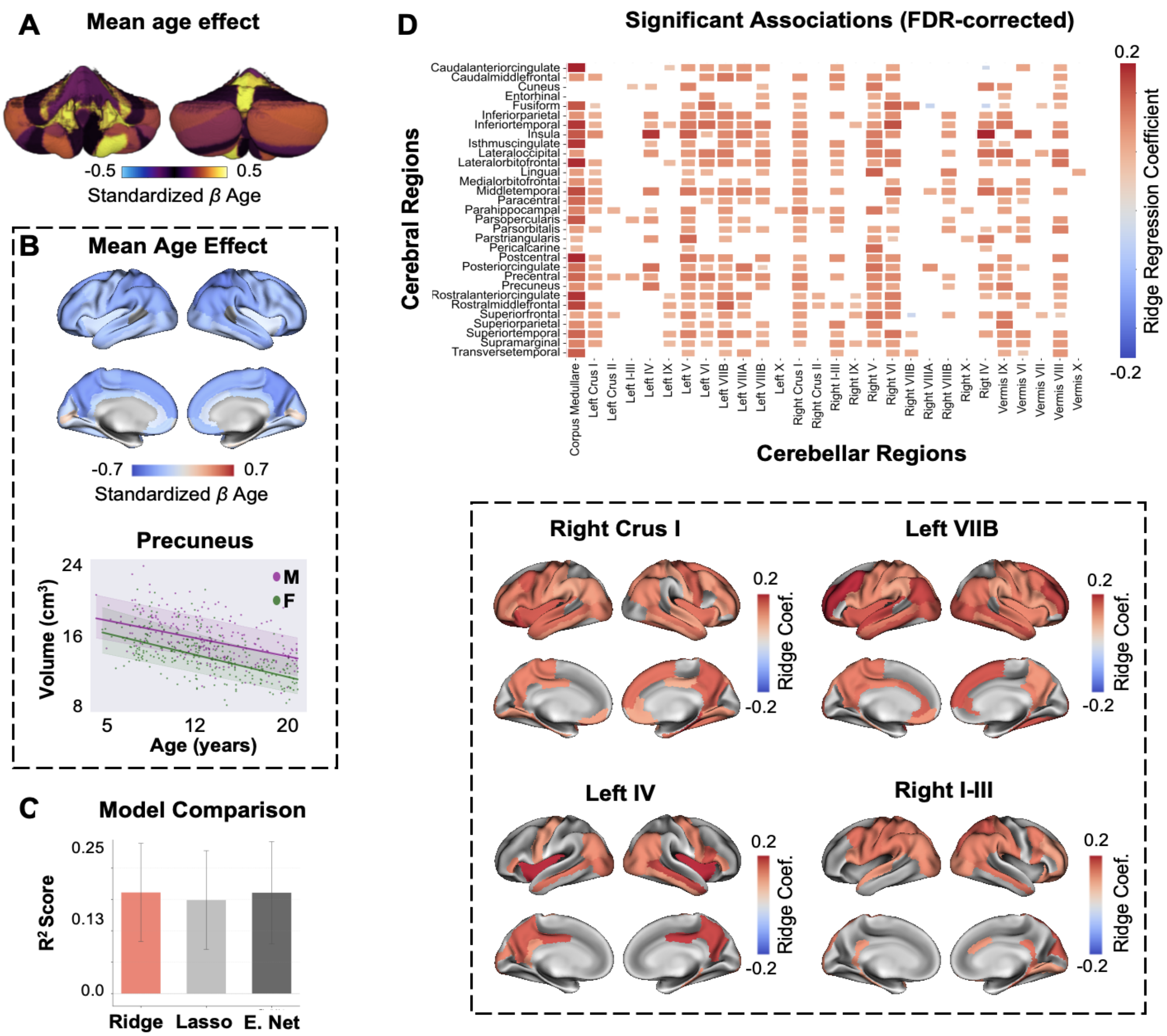

**Supplementary Figure 11.** Association of cerebellar and cerebral growth trajectories. **A.** Mean effect of age on growth per anatomical lobule (HCP-D dataset; 5-21 years). **B.** Mean effect of age on growth per DK parcel (top) and an example normative trajectory for the precuneus (bottom). Bold lines represent the mean trajectory per sex. Shaded areas represent the 95.5% confidence interval. **C.** Comparison of Ridge, Lasso, and ElasticNet regularization models based on $R^2$ scores (ten-fold cross-validation). Error bars represent the standard error of the mean. **D.** Top: Significant cerebro-cerebellar associations (10,000 permutations, false discovery rate (FDR)-corrected at $q = .05$). Left and right DK parcels in the heatmap are averaged for brevity. Bottom: FDR-corrected weights for example anterior (lobules I-III and IV) and posterior (Right Crus I and Left VIIB) cerebellar parcels, projected on the DK atlas. Abbreviations: Coef. = coefficient; M = male; F = female; E. Net = ElasticNet.

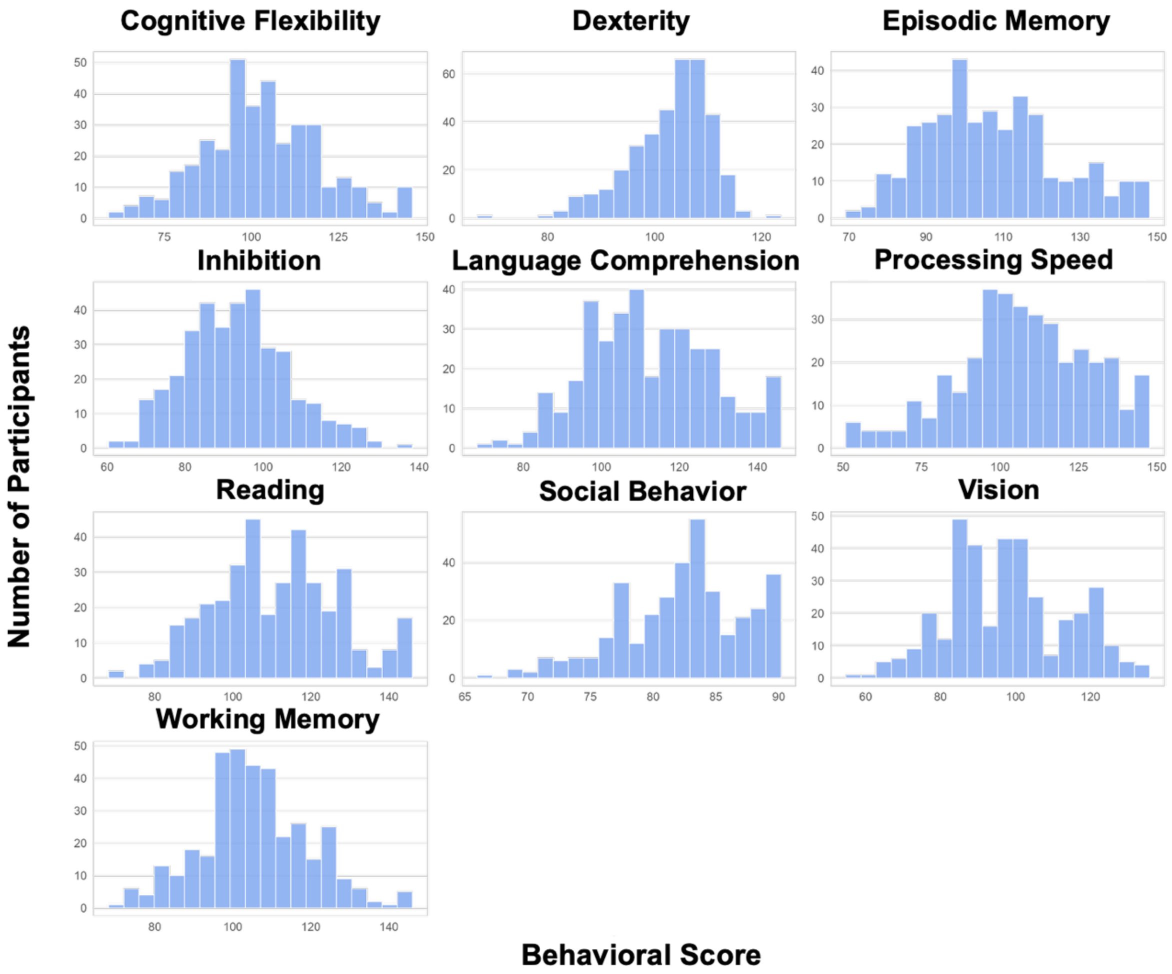

**Supplementary Figure 12.** Distribution of participants' behavioral scores in the Human Connectome Project Development dataset (*N* = 457; 5-21 years; participants without behavioral scores were removed).

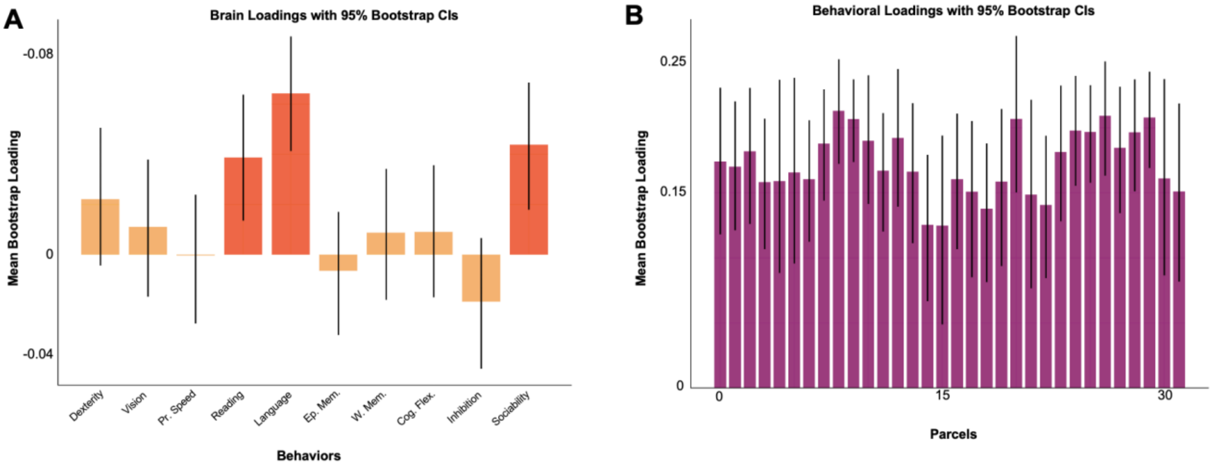

**Supplementary Figure 13.** Bootstrap resampling of PLS loadings. Bootstrap resampling (10,000 iterations) results for behavioral scores **(A)** and functional fusion parcels **(B)**. Bars correspond to 95% confidence intervals (CIs). Abbreviations: Pr. Speed = Processing Speed; Ep. Mem. = Episodic Memory; W. Mem. = Working Memory; Cog. Flex. = Cognitive Flexibility.

**Supplementary Table 1. Sex-stratified means of standardized $\beta_{age}$ across lobular regions**

| Lobular Regions | Male (Mean [95% CI]) | Female (Mean [95% CI]) |
|---|---|---|
| Corpus Medullare | 0.46 [-0.08 1.04] | 0.98 [0.26 1.84] |
| Left Crus I | 0.40 [-0.22 0.91] | 0.34 [-0.21 0.99] |
| Left Crus II | 0.36 [-0.15 0.97] | 0.35 [-0.27 1.38] |
| Left I-III | 0.03 [-0.22 0.25] | 0.02 [-0.54 0.63] |
| Left IV | 0.40 [-0.15 0.80] | 0.32 [-0.12 0.82] |
| Left IX | 0.04 [-0.64 0.67] | 0.42 [-0.27 1.24] |
| Left V | 0.41 [-0.22 0.89] | 0.07[-0.13 0.87] |
| Left VI | 0.42 [-0.23 1.09] | 0.26 [-0.37 0.91] |
| Left VIIB | 0.61 [0.01 1.17] | 0.65 [0.34 1.87] |
| Left VIIIA | 0.08 [-0.42 0.59] | -0.08 [-0.54 0.36] |
| Left VIIIB | 0.40 [-0.02 0.85] | 0.20 [-0.34 0.72] |
| Left X | 0.05 [-0.45 0.51] | -0.05 [-0.47 0.36] |
| Right Crus I | 0.48 [-0.10 1.02] | 0.45 [-0.17 1.10] |
| Right Crus II | 0.40 [-0.14 1.23] | 0.34 [-0.20 1.11] |
| Right I-III | 0.26 [-0.20 0.81] | 0.29 [-0.09 1.89] |
| Right IX | 0.13 [-0.49 0.78] | 0.32 [-0.17 1.65] |
| Right V | 0.49 [0.06 0.91] | 0.41 [-0.08 0.93] |
| Right VI | 0.32 [-0.27 0.86] | 0.62 [-0.05 2.01] |
| Right VIIB | 0.78 [0.17 1.48] | 0.79 [0.29 1.97] |
| Right VIIIA | 0.05 [-0.60 0.57] | 0.05 [-0.40 0.64] |
| Right VIIIB | 0.29 [-0.19 0.76] | 0.32 [-0.24 0.87] |
| Right X | -0.02 [-0.52 0.48] | -0.08 [-0.43 0.29] |
| Right IV | 0.25 [-0.15 0.66] | 0.07 [-0.30 0.44] |
| Vermis IX | 0.28 [-0.17 0.72] | 0.25 [-0.27 0.87] |
| Vermis VI | 0.34 [-0.15 0.78] | 0.34 [0.02 1.12] |
| Vermis VII | 0.22 [-0.05 0.47] | 0.06 [-0.55 0.38] |
| Vermis VIII | 0.06 [-0.47 0.56] | 0.05 [-0.42 0.52] |
| Vermis X | 0.05 [-0.16 0.26] | 0.03 [-0.25 0.27] |

Abbreviations: CI = confidence interval

**Supplementary Table 2. Sex-stratified means of standardized $\beta_{age}$ across fusion regions.**

| Functional Fusion Regions | Male (Mean [95% CI]) | Female (Mean [95% CI]) |
|---|---|---|
| A1L | 0.38 [0.25 0.49] | 0.3 [0.21 0.38] |
| A1R | 0.36 [0.24 0.49] | 0.32 [0.12 0.32] |
| A2L | 0.3 [0.18 0.43] | 0.16 [0.06 0.25] |
| A2R | 0.22 [0.1 0.34] | 0.14 [0.04 0.24] |
| A3L | 0.33 [0.22 0.45] | 0.21 [0.12 0.29] |
| A3R | 0.28 [0.17 0.4] | 0.2 [0.11 0.31] |
| D1L | 0.53 [0.41 0.64] | 0.49 [0.26 0.46] |
| D1R | 0.46 [0.34 0.58] | 0.34 [0.14 0.35] |
| D2L | 0.35 [0.23 0.46] | -0.12 [-0.20 0.33] |
| D2R | 0.38 [0.26 0.48] | 0.18 [0.08 0.28] |
| D3L | 0.23 [0.11 0.36] | 0.13 [0.02 0.23] |
| D3R | 0.43 [0.32 0.55] | 0.27 [0.17 0.37] |
| D4L | 0.23 [0.11 0.36] | 0.14 [0.04 0.23] |
| D4R | 0.48 [0.38 0.59] | 0.3 [0.2 0.4] |
| M1L | 0.45 [0.34 0.55] | 0.28 [0.18 0.37] |
| M1R | 0.39 [0.28 0.52] | 0.28 [0.17 0.38] |
| M2L | -0.21 [-0.32 0.45] | 0.12 [0.02 0.22] |
| M2R | 0.22 [0.1 0.35] | -0.11 [-0.20 0.22] |
| M3L | 0.4 [0.28 0.51] | 0.20 [0.08 0.28] |
| M3R | 0.13 [0 0.24] | 0.03 [-0.07 0.14] |
| M4L | 0.03 [-0.05 0.41] | 0.13 [0.04 0.22] |
| M4R | 0.16 [0.04 0.28] | 0.08 [-0.02 0.18] |
| S1L | 0.53 [0.42 0.63] | 0.31 [0.22 0.4] |
| S1R | 0.4 [0.27 0.51] | 0.18 [0.07 0.28] |
| S2L | 0.41 [0.39 0.62] | 0.54 [0.24 0.45] |
| S2R | 0.29 [0.19 0.41] | 0.09 [0 0.19] |
| S3L | 0.46 [0.34 0.58] | -0.08 [-0.18 0.38] |
| S3R | 0.28 [0.17 0.4] | 0.11 [0.01 0.22] |
| S4L | 0.32 [0.2 0.43] | 0.17 [0.07 0.27] |
| S4R | 0.3 [0.19 0.41] | -0.17 [-0.07 0.27] |
| S5L | 0.11 [-0.01 0.22] | 0.04 [-0.06 0.14] |
| S5R | 0.17 [0.04 0.28] | 0.1 [0.01 0.2] |

Abbreviations: CI = confidence interval; A = Action; D = Demand; M = Motor; S = Socio-Linguistic; L = left; R = right.

**Supplementary Table 3. Sex-stratified means of standardized $\beta_{age}$ across MDTB regions.**

| MDTB Regions | Male (Mean [95% CI]) | Female (Mean [95% CI]) |
|---|---|---|
| 1 (Left-Hand Presses) | 0.31 [0.18 0.44] | 0.11 [0.02 0.22] |
| 2 (Right-Hand Presses) | 0.16 [0.04 0.29] | 0.08 [-0.02 0.18] |
| 3 (Saccades) | 0.31 [0.16 0.41] | 0.26 [0.16 0.35] |
| 4 (Action Observation) | 0.31 [0.19 0.43] | 0.23 [0.13 0.33] |
| 5 (Divided Attention Left) | 0.42 [0.29 0.54] | 0.3 [0.19 0.4] |
| 6 (Divided Attention Right) | 0.19 [0.11 0.4] | 0.12 [0.09 0.32] |
| 7 (Narrative) | 0.49 [0.38 0.59] | 0.34 [0.25 0.43] |
| 8 (Word Comprehension) | 0.26 [0.15 0.38] | 0.18 [0.08 0.27] |
| 9 (Verbal Fluency) | 0.27 [0.15 0.38] | 0.16 [0.05 0.26] |
| 10 (Autobiographical Recall) | 0.39 [0.23 0.46] | 0.3 [0.2 0.4] |

Abbreviations: CI = confidence interval.

**Supplementary Table 4: Sex-stratified means of standardized $\beta_{age}$ across resting-state regions.**

| Resting-state regions | Male (Mean [95% CI]) | Female (Mean [95% CI]) |
|---|---|---|
| Somatomotor | 0.15 [0.10 0.39] | 0.21 [0.11 0.31] |
| Dorsal Attention | 0.39 [0.27 0.5] | 0.26 [0.16 0.36] |
| Ventral Attention | 0.4 [0.28 0.51] | 0.25 [0.16 0.35] |
| Limbic | 0.12 [0.09 0.32] | 0.22 [0.02 0.22] |
| Frontoparietal | 0.23 [0.32 0.55] | 0.35 [0.24 0.44] |
| Default | 0.37 [0.25 0.47] | 0.22 [0.15 0.35] |

Abbreviations: CI = confidence interval.

**Supplementary Table 5. Lobular region model comparison via leave-one-out cross-validation (LOOCV).**

| Lobular Regions | Linear Model LOOCV [SE] | B-Spline Model LOOCV [SE] | Difference [SE of difference] |
|---|---|---|---|
| Corpus Medullare | -665.82 [22.38] | -647.33 [21.04] | 18.49 [4.64] |
| Left Crus I | -770.53 [22.63] | -766.98 [22.15] | 3.54 [2.44] |
| Left Crus II | -794.2 [18.42] | -790.58 [18.2] | 3.62 [2.98] |
| Left I-III | -115.1 [32.82] | -116 [32.55] | 0.9 [1.89] |
| Left IV | -750.41 [19.69] | -747.2 [19.41] | 3.21 [2.48] |
| Left IX | -809.4 [18.09] | -807.33 [18] | 2.08 [2.22] |
| Left V | -727.11 [17.86] | -725.86 [17.93] | 1.25 [2.96] |
| Left VI | -769.62 [21.49] | -769.88 [21.45] | 0.26 [2.12] |
| Left VIIB | -721.83 [18.22] | -713.76 [18.53] | 8.07 [4.36] |
| Left VIIIA | -795.63 [19.68] | -795.98 [19.61] | 0.35 [1.87] |
| Left VIIIB | -762.07 [19.78] | -763.21 [19.87] | 1.14 [1.41] |
| Left X | -828 [20.13] | -830.26 [20.27] | 2.26 [1.35] |
| Right Crus I | -761.26 [19.76] | -758.19 [19.62] | 3.06 [2.47] |
| Right Crus II | -788.1 [18.68] | -780.34 [18.41] | 7.76 [3.79] |
| Right I-III | -810.84 [17.4] | -808.88 [17.62] | 1.96 [2.4] |
| Right IX | -814.71 [16.79] | -813.08 [16.74] | 1.63 [2.18] |
| Right V | -716.06 [28.06] | -709.08 [25.54] | 6.98 [4.96] |
| Right VI | -748.76 [16.3] | -741.88 [16.22] | 6.88 [3.48] |
| Right VIIB | -753.64 [17.05] | -748.1 [17.52] | 5.54 [3.78] |
| Right VIIIA | -795.09 [19.56] | -792.47 [19.3] | 2.62 [2.49] |
| Right VIIIB | -753.95 [18.67] | -753.88 [18.82] | 0.07 [2.03] |
| Right X | -821.77 [20.95] | -821.48 [20.52] | 0.28 [2.35] |
| Right IV | -781.38 [17.98] | -784.03 [18.04] | 2.65 [1.05] |
| Vermis IX | -774.66 [26.45] | -770.02 [25.82] | 4.64 [2.96] |
| Vermis VI | -728.81 [23.38] | -729.05 [23.24] | 0.24 [2.34] |
| Vermis VII | -333.64 [33.03] | -333.24 [32.98] | 0.4 [2.35] |
| Vermis VIII | -697.9 [24.19] | -696.18 [23.75] | 1.72 [2.96] |
| Vermis X | -230.5 [33.09] | -228.2 [33.58] | 2.29 [3.03] |

Abbreviations: SE = standard error of the mean.

**Supplementary Table 6. Functional fusion region model comparison via leave-one-out cross-validation (LOOCV).**

| Fusion Regions | Linear Model LOOCV [SE] | B-Spline Model LOOCV [SE] | Difference [SE of difference] |
|---|---|---|---|
| A1L | -646.63 [17.54] | -643.58 [17.6] | 3.05 [3.35] |
| A1R | -685.45 [15.42] | -676.76 [15.74] | 8.69 [4.97] |
| A2L | -663.33 [16.53] | -664.58 [16.64] | 1.25 [2.02] |
| A2R | -693.69 [15.3] | -694.01 [15.71] | 0.31 [1.67] |
| A3L | -645.26 [14.35] | -644.26 [14.52] | 1 [2.7] |
| A3R | -701.75 [14.48] | -701.75 [14.48] | 0 [1.87] |
| D1L | -649.23 [19.5] | -643.6 [20.27] | 5.63 [4.24] |
| D1R | -674.56 [15.71] | -671.79 [15.65] | 2.77 [3.42] |
| D2L | -660.23 [16.22] | -653.58 [16.4] | 6.64 [4.16] |
| D2R | -660.6 [14.82] | -656.82 [15.17] | 3.78 [3.87] |
| D3L | -693.3 [16.9] | -690.37 [17.24] | 2.93 [3.3] |
| D3R | -667.2 [16.2] | -666.57 [16.51] | 0.63 [2.63] |
| D4L | -682.59 [16.6] | -677.96 [16.67] | 4.62 [3.36] |
| D4R | -660.46 [16.96] | -657.04 [17.44] | 3.42 [3.21] |
| M1L | -639.76 [22.72] | -640.66 [22.62] | 0.9 [2.05] |
| M1R | -680.67 [18.38] | -682.56 [18.29] | 1.89 [1.73] |
| M2L | -681.22 [19.73] | -682.17 [19.5] | 0.95 [1.91] |
| M2R | -701.76 [16.8] | -704.09 [16.85] | 2.33 [1.32] |
| M3L | -678.22 [17.41] | -679.61 [17.6] | 1.38 [2.19] |
| M3R | -715.15 [15.71] | -715.29 [15.58] | 0.14 [2.3] |
| M4L | -650.37 [17.59] | -650.39 [17.53] | 0.02 [2.41] |
| M4R | -695.99 [15.25] | -696.44 [15.17] | 0.45 [1.7] |
| S1L | -611.45 [19.78] | -612.08 [19.58] | 0.63 [2.49] |
| S1R | -691.66 [14.25] | -683.91 [14.11] | 7.75 [4.88] |
| S2L | -650.93 [14.84] | -641.39 [14.96] | 9.54 [4.9] |
| S2R | -685.85 [14.3] | -682.35 [15.16] | 3.5 [3.75] |
| S3L | -653.88 [15.28] | -645.04 [15.1] | 8.84 [5.07] |
| S3R | -689.04 [15.17] | -689.23 [15.27] | 0.2 [2.29] |
| S4L | -678.55 [16.06] | -674.06 [16.34] | 4.49 [3.59] |
| S4R | -653.07 [13.86] | -650.33 [14.36] | 2.74 [3.53] |
| S5L | -694.11 [16.98] | -692.16 [16.5] | 1.95 [3.14] |
| S5R | -701.69 [16.03] | -703.34 [15.97] | 1.65 [1.15] |

Abbreviations: SE = standard error of the mean; A = Action; D = Demand; M = Motor; S = Socio-Linguistic; L = left; R = right.

**Supplementary Table 7. Functional MDTB region model comparison via leave-one-out cross-validation (LOOCV).**

| MDTB Regions | Linear Model LOOCV [SE] | B-Spline Model LOOCV [SE] | Difference [SE of difference] |
|---|---|---|---|
| 1 (Left-Hand Presses) | -669.67 [15.65] | -672.24 [15.48] | 2.57 [2.17] |
| 2 (Right-Hand Presses) | -676.46 [14.21] | -678.76 [14.61] | 2.31 [1.55] |
| 3 (Saccades) | -662.75 [15.88] | -661.68 [15.85] | 1.07 [2.59] |
| 4 (Action Observation) | -646.35 [15.16] | -641.6 [15.85] | 4.75 [4.4] |
| 5 (Divided Attention Left) | -656.98 [15.71] | -651.02 [15.82] | 5.96 [4.24] |
| 6 (Divided Attention Right) | -661.2 [15.04] | -660.45 [15.64] | 0.74 [2.74] |
| 7 (Narrative) | -628.46 [18.35] | -627.62 [18] | 0.84 [3.27] |
| 8 (Word Comprehension) | -649.23 [13.42] | -645.02 [13.88] | 4.2 [3.69] |
| 9 (Verbal Fluency) | -666.14 [14.44] | -666.13 [14.82] | 0.01 [2.5] |
| 10 (Autobiographical Recall) | -641.19 [16.01] | -631.63 [16.15] | 9.57 [4.74] |

Abbreviations: SE = standard error of the mean.

**Supplementary Table 8. Functional resting-state region model comparison via leave-one-out cross-validation (LOOCV).**

| Resting-State Regions | Linear Model LOOCV [SE] | B-Spline Model LOOCV [SE] | Difference [SE of difference] |
|---|---|---|---|
| Somatomotor | -667.48 [15.23] | -666.07 [15.06] | 1.41 [2.69] |
| Dorsal Attention | -652.87 [14.61] | -639.9 [14.4] | 12.97 [5.46] |
| Ventral Attention | -641.69 [14.1] | -634.59 [13.43] | 7.1 [4.61] |
| Limbic | -674.66 [17.06] | -667.8 [16.71] | 6.86 [4.53] |
| Frontoparietal | -648.15 [15.61] | -639.38 [15.72] | 8.77 [4.66] |
| Default | -646.72 [14.06] | -633.08 [14.26] | 13.64 [5.84] |

Abbreviations: SE = standard error of the mean.

**Supplementary Table 9. Significant PLS bootstrap ratios.**

| Variable Type | Variable | BSR | 95% CI |
|---|---|---|---|
| *Behavior* | Language Comprehension | 5.50 | [0.04 0.09] |
| | Social Behavior | 3.44 | [0.02 0.07] |
| | Reading | 3.18 | [0.02 0.06] |
| *Parcel* | D2R | 12.52 | [0.17 0.24] |
| | D2L | 10.38 | [0.17 0.26] |
| | S4R | 10.33 | [0.17 0.25] |
| | S2R | 10.02 | [0.15 0.23] |
| | S3L | 8.83 | [0.16 0.25] |
| | S4L | 8.74 | [0.15 0.24] |
| | S2L | 8.71 | [0.15 0.24] |
| | D1R | 8.15 | [0.14 0.23] |
| | S3R | 7.73 | [0.14 0.23] |
| | D3L | 7.43 | [0.14 0.24] |
| | D3R | 7.23 | [0.12 0.21] |
| | A2L | 7.04 | [0.13 0.24] |
| | D4L | 7.01 | [0.14 0.24] |
| | M4L | 6.99 | [0.16 0.27] |
| | A1R | 6.74 | [0.12 0.23] |
| | S1R | 6.59 | [0.12 0.23] |
| | A2R | 6.34 | [0.11 0.21] |
| | D1L | 6.15 | [0.11 0.20] |
| | A1L | 5.93 | [0.12 0.23] |
| | M2L | 5.81 | [0.10 0.21] |
| | D4R | 5.52 | [0.10 0.22] |
| | M3R | 5.22 | [0.10 0.22] |
| | M3L | 5.19 | [0.09 0.19] |
| | M2R | 4.90 | [0.09 0.21] |
| | A3R | 4.88 | [0.10 0.24] |
| | S5R | 4.71 | [0.09 0.22] |
| | S5L | 4.55 | [0.09 0.24] |
| | S1L | 4.45 | [0.08 0.20] |
| | A3L | 4.28 | [0.09 0.23] |
| | M1L | 4.03 | [0.05 0.18] |
| | M4R | 3.73 | [0.08 0.23] |

| Variable Type | Variable | BSR | 95% CI |
|---|---|---|---|
| | M1R | 3.45 | [0.05 0.19] |

Only variables with |BSR| > 2 and 95% CI not crossing zero are shown. Variables are ordered by bootstrap ratio magnitude within each type. Abbreviations: CI = confidence interval; BSR = bootstrap ratio.
